# Acetyl-CoA Carboxylase Obstructs CD8^+^ T-Cell Lipid Utilization and Energy Synthesis in the Tumor Microenvironment

**DOI:** 10.1101/2023.11.16.567439

**Authors:** Elizabeth G. Hunt, Katie E. Hurst, Andrew S. Kennedy, Brian P. Riesenberg, Evelyn J. Gandy, Alex M. Andrews, Peng Gao, Michael F. Coleman, Emily D. Wallace, Jessica E. Thaxton

## Abstract

The solid tumor microenvironment (TME) imprints a compromised metabolic state in tumor infiltrating T cells (TILs) hallmarked by the inability to maintain effective energy synthesis for antitumor function and survival. T cells in the TME must catabolize lipids via mitochondrial fatty acid oxidation (FAO) to supply energy in nutrient stress, and it is established that T cells enriched in FAO are adept at cancer control. However, endogenous TILs and unmodified cellular therapy products fail to sustain bioenergetics in tumors. Using patient samples and mouse models, we reveal that the solid TME imposes perpetual acetyl-CoA carboxylase (ACC) activity, enforcing lipid biogenesis and storage in TILs that directly opposes FAO. Using metabolic, lipidomic, and confocal imaging strategies, we find that restricting ACC wholly rewires T cell metabolism, enabling energy maintenance in TME stress. Moreover, limiting ACC activity potentiates a gene and phenotypic program indicative of T cell memory, engendering TILs with increased survival and polyfunctionality, with the ability to control solid cancer.

## Introduction

T cells are a subset of adaptive immunity with robust cytotoxic capability that promote control of cancers. In the solid TME, T cells experience loss of function, limiting cancer control. A reason for loss of T cell efficacy is that the solid TME contains limited nutrient resources for which tumor cells and immune cells must compete to fuel cellular functions (Kouidhi et al., 2017). Upon entering the TME, T cell populations experience extensive stress and inconsistent nutrient stress supply, prompting translation attenuation and metabolic dysfunction (Rial Saborido et al., 2022; Riesenberg et al., 2022). Specifically, there is a competition for glucose disproportionately won by tumor cells, leading to intermittent pockets of glucose availability in the TME (Chang et al., 2015; Reinfeld et al., 2021). Cell-intrinsic programs, largely controlled by mTORC1 signaling, contribute to this uneven division of nutrients between tumor cells and various immune cells (Reinfeld *et al*., 2021). Because cytotoxic T cells rely on aerobic glycolysis to produce cytokines such as IFN-γ (MacIver et al., 2013), tumor-mediated glucose restriction can impair T cell antitumor effector function (Chang *et al*., 2015). Taken together, strategies that alter T cell metabolic dependencies and diminish reliance on the exogenous microenvironment could augment cellular energetics to enhance the efficacy of T cell-based immunotherapies in solid cancers.

In non-immune cell types, it is established that fasted states force a shift in cellular metabolism from utilization of glycolysis toward oxidation of free fatty acids (FAO) within the mitochondria to sustain energy synthesis in nutrient stress (Hardie, 2003). Preclinical and clinical evidences show that T cell products enriched in FAO enable prolonged and superior T cell antitumor function (Klebanoff et al., 2005; Xu et al., 2014) (van der Windt et al., 2012; Windt et al., 2013), but the majority of endogenous tumor infiltrating T cells appear restricted in the capacity to employ FAO (Chamoto et al., 2017; Scharping et al., 2016). FAO is initiated by the conversion of energy-rich fatty acyl-CoA to acyl-carnitine at the mitochondrial outer membrane by the enzyme carnitine palmitoyltransferase 1A (CPT1A) followed by transport across the inner mitochondrial membrane and subsequent conversion back to fatty acyl-CoA (Lee et al., 2011). Available fatty acyl-CoA in the mitochondrial matrix enables cycles of fatty acid β-oxidation, producing acetyl-CoA that can enter the TCA, providing NADH and FADH2 equivalents utilized by the electron transport chain to generate ATP (Martínez-Reyes and Chandel, 2020). In fasting states, cells utilize FAO to sustain ATP synthesis. Thus, in T cell cultures, limiting glucose availability by targeting Akt (Crompton et al., 2015) or by directly restricting glycolysis (Sukumar et al., 2013) results in a pool of T cells with heightened dependence on FAO. Infusion of these cell products into tumor-bearing hosts augments T cell performance against solid tumors, establishing that altering T cell metabolism is a strategy to augment T cell products for clinical infusion. However, it remains poorly understood how lipid biology is regulated in CD8^+^ T cells in the solid TME, and the molecular players that restrict utilization of FAO-based metabolism in TILs.

Cytosolic fatty acid synthesis (FAS) directly opposes mitochondrial fat oxidation and is governed by the enzyme acetyl-CoA carboxylase (ACC). ACC induces carboxylation of carbohydrate-derived acetyl-CoA in the cytosol, forming malonyl-CoA which is the metabolic precursor to fatty acids. Thus, through formation of malonyl-CoA, ACC orchestrates the first committed step in FAS. Two isoforms of the ACC enzyme are known, ACC1 and ACC2, each encoded by different genes and differentially expressed based on tissue and cell type (Wakil and Abu-Elheiga, 2009). ACC1 is predominantly expressed in lipogenic tissues including liver, lactating mammary gland, and adipose. ACC1-generated malonyl-CoA is chiefly used by fatty acid synthase (FASN) to generate fatty acids in the cytosol. Moreover, through reducing the intracellular supply of acetyl-CoA, an important cofactor for histone acetylation enzymes, ACC1 activity influences transcriptional regulation in the nucleus (Galdieri and Vancura, 2012). ACC2 is principally expressed in muscle and heart (Abu-Elheiga et al., 1997), localizing to the mitochondrial outer membrane where ACC2-generated malonyl-CoA potently restricts FAO in the mitochondria through inhibition of CPT1A (Hardie and Pan, 2002; Wakil and Abu-Elheiga, 2009). In eight human non-small-cell lung cancer (NSCLC) cell lines, gene deletion of ACC1 was sufficient to reduce NSCLC viability *in vitro* and impair tumorigenesis *in vivo,* highlighting the requirement of ACC-regulated FAS for tumor cell growth (Svensson et al., 2016). The contribution of ACC to shape CD8^+^ TIL metabolism is not known. In CD4^+^ T cells, gene deletion of ACC1 generated T cell memory with heightened FAO-dependence indicating that limiting ACC was sufficient to program T cell metabolic and lineage fate (Endo et al., 2019). However, CD8^+^ T cell programming through ACC has not been addressed, nor has ACC been studied as a target that could restrict effective T cell bioenergetics in solid tumors.

Here we hypothesized that ACC expression in CD8^+^ TILs could impede metabolic plasticity of CD8^+^ TILs, restricting utilization of FAO to support sustained bioenergetics in solid cancers. Using RNA-sequencing (RNA-seq) paired with a proteomic screen, we observed that ACC1 is induced in mouse and human CD8^+^ TILs coincident with a significant increase in steatosis evidenced by accumulation of lipid droplets (LDs). Using metabolomics, we found that ACC inhibition rewired CD8^+^ T cell metabolism, resulting in a shift toward FAO utilization. Inhibition of ACC in CD8^+^ T cells exposed to chronic nutrient stress resulted in accelerated ATP synthesis relative to vehicle controls, and the effect was dependent on CPT1A-mediated lipid import. We uncovered that ACC directly promotes storage of triacylglycerides (TAGs) in cytosolic LDs, prohibiting the utilization of free fatty acids (FFA) by the mitochondria. When ACC activity was restricted, heightened activity within the endoplasmic reticulum (ER)-mediated TAG synthesis pathway resulted in immediate utilization of FFAs by mitochondria for FAO-mediated energy maintenance. Unexpectedly, inhibition of ACC in mouse and human T cells produced T cell pools endowed with phenotypic traits associated with T cell stemness and longevity. CD8^+^ T cells conditioned with an ACC inhibitor engendered long-term control of melanomas marked by *in vivo* persistence and increased polyfunctionality.

## Results

### ACC1 is enriched in CD8^+^ TILs coincident with steatosis

Poor metabolic function, such as diminished mitochondrial ATP stores, has been linked to the inefficacy of T cells in solid cancers (Chamoto *et al*., 2017; Scharping *et al*., 2016). We performed RNA-seq on CD8^+^ TILs isolated from MCA-205 fibrosarcomas and autologous splenic CD8^+^ T cells, then probed the data set for gene signatures associated with poor metabolic prognosis. Gene set enrichment analysis (GSEA) identified that CD8^+^ TILs were significantly enriched for genes associated with abnormal circulating lipid relative to autologous splenocytes (Figure 1a), suggestive that irregular lipid accumulation could afflict CD8^+^ T cells in tumors. The data prompted us to perform untargeted ultra-performance liquid chromatography coupled to mass spectrometry (UPLC-MS)-based lipidomics on CD8^+^ TILs isolated from MCA-205 fibrosarcomas and autologous splenic CD8^+^ T cells. Consistent with the RNA-seq data set, integration of peak area of all lipids identified suggested that CD8^+^ TILs harbored ∼1.5-fold increase in abundance of lipids compared to CD8^+^ T cells isolated from spleens (Figure 1b). Next, we analyzed the data set through identification of individual lipid classes to determine whether a particular group of lipids was preferentially enriched in CD8^+^ TILs. We observed that the majority of lipid classes including triacylglycerides (TAG), diacylglycerides (DAG), phosphatidylcholines (PC), phosphatidylglycans (PG), phosphatidylinositols (PI), and phosphatidylethanolamines (PE) were significantly increased in CD8^+^ TILs compared to CD8^+^ T cells isolated from spleens (Figure 1c). Together, the data indicate that T cells in tumors harbor increased lipids relative to peripheral CD8^+^ T cells.

**Figure 1.**
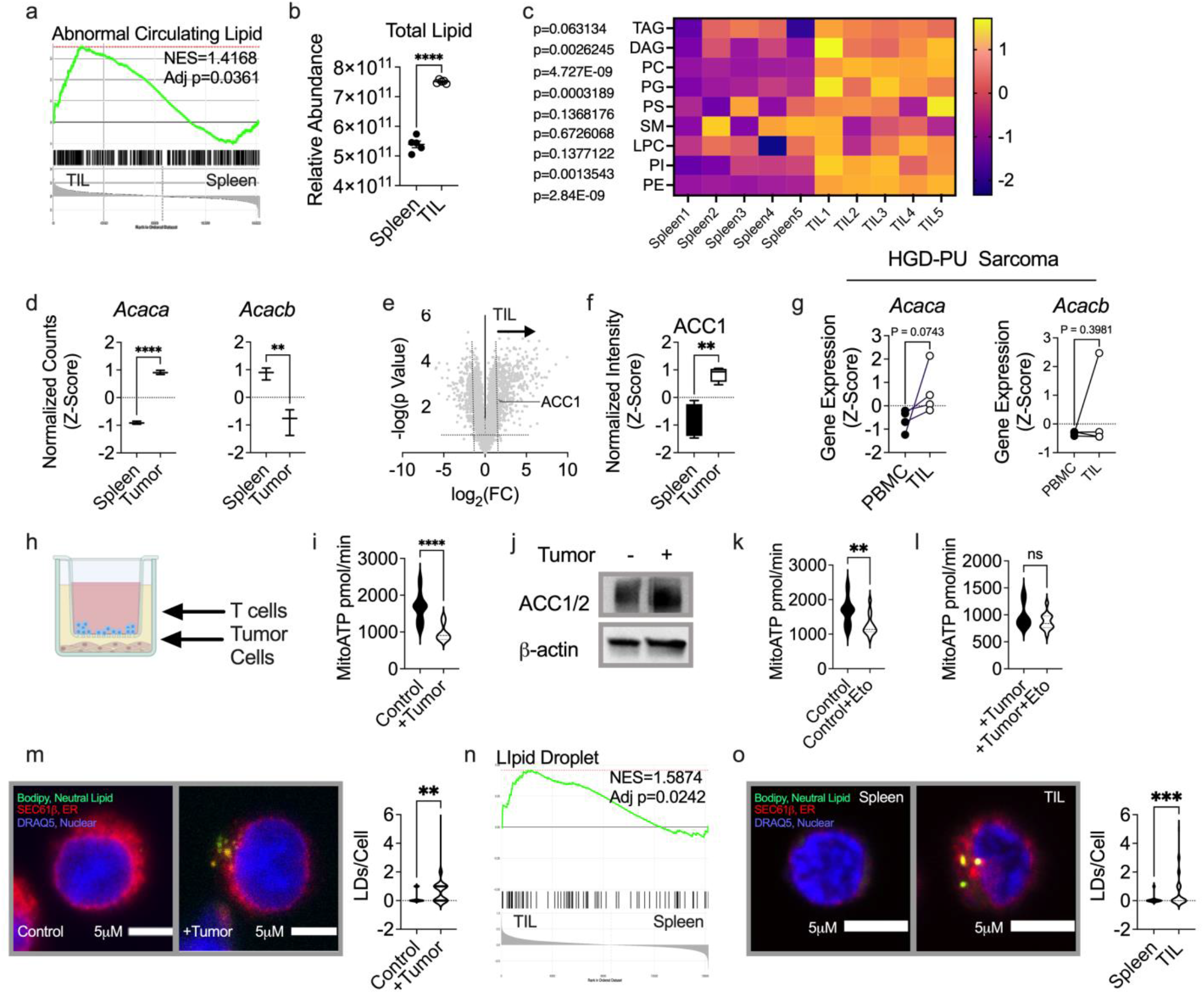
ACC1 is enriched in CD8^+^ TILs coincident with steatosis. CD8^+^ T cells were FACS sorted from spleens and tumors of C57BL/6 mice bearing MCA-205 fibrosarcomas. RNA-sequencing was performed on CD8^+^ T cells pooled from n=3 groups of mice. a) Leading edge plot generated from GSEA of RNA-sequencing data. UPLC-MS was performed on CD8^+^ T cells pooled from n=5 groups of mice. b) Quantification of integration of total peak area of lipids detected and c) relative abundance of individual lipid classes identified between CD8^+^ splenocytes and TILs. d) Normalized reads of indicated genes from RNA-sequencing data set described in a. Proteomics was performed on CD8^+^ T cells pooled from n=5 groups of mice. e) Volcano plot of proteins identified by the proteomics screen in CD8^+^ splenocytes (left) and TILs (right) and f) normalized intensity of ACC1 protein is shown. CD8^+^ T cells were bead sorted from autologous patient peripheral blood and tumor tissue and g) RT-PCR for indicated gene expression was performed (n=4 patients). h) Schematic of tumor cell-T cell transwell co-culture assay. OT-1 T cells were activated with OVA peptide in the presence of IL-2 and expanded for 3 days then seeded into control or transwells seeded with B16F1 melanoma cells. After 36 hours of co-culture, T cells were collected and Seahorse Real-Time ATP Rate Assay was performed. Quantifications among T cell groups of i, k-l) mitochondrial ATP rates (mitoATP) are shown in response to sequential injections of i, oligomycin and rotenone/antimycin A or k-l, etomoxir or buffer control, oligomycin, and rotenone/antimycin A. mitoATP rate was calculated based on the decrease in the OCR. Data points represent individual Seahorse wells. T cells were co-cultured as in h, then collected for j) western blotting or m) confocal imaging using Bodipy 493/503 to visualize LDs. Data points represent number of LDs per cell (n=50). n) Leading edge plot generated from GSEA of RNA-sequencing data set described in a. CD8^+^ T cells were FACS sorted from spleens and tumors of C57BL/6 mice bearing MCA-205 fibrosarcomas and o) confocal imaging using Bodipy 493/503 to visualize LDs was performed, (n=50 cells quantified). Seahorse, western blotting, and imaging independent experiments repeated at least 3 times. ns, not significant, * p < 0.05, ** p < 0.01, *** p < 0.001, **** p < 0.0001 based on statistical analysis by unpaired or paired (g) Student’s t-test, all error bars indicate the SEM.

The enzyme ACC controls *de novo* lipogenesis and has been targeted for non-alcoholic fatty liver disease (NAFLD) due to its role in promotion of excess lipids and impairment of FAO (Imai and Cohen, 2018). Given the robust induction of lipids in CD8^+^ TILs relative to peripheral T cells, we probed the RNA-seq data set for *Acaca* (ACC1) and *Acacb* (ACC2) isoenzymes. Normalized reads indicated that *Acaca* expression was significantly increased in CD8^+^ TILs, but *Acacb* expression was reduced relative to splenocytes (Figure 1d). Next, we performed a proteomics screen to identify whether ACC1 or ACC2 could be detected at the protein level. We observed that ACC1 was significantly upregulated in CD8^+^ TILs relative to T cells isolated from spleens (Figure 1e-f). ACC2 was undetected in T cells isolated from both sites, possibly indicative that protein expression of ACC2 was below the level of instrument detection. To determine whether the enzymes were regulated at the gene level in cancer patients we obtained fresh untreated tumor tissue and matched peripheral blood from pleiomorphic undifferentiated high-grade deep (PU HGD) sarcoma patients and measured gene expression using real-timePCR. Similar to the sarcoma mouse model, we observed that *Acaca* expression was increased in CD8^+^ patient TILs relative to peripheral CD8^+^ T cells, but *Acacb* did not show a similar trend (Figure 1g). These data suggest that ACC1 is induced in T cells in primary sarcomas.

To understand the functional role that ACC1/2 could play in regulation of metabolic efficacy of CD8^+^ TILs, we utilized an *in vitro* transwell assay in which tumor cells and T cells are co-cultured for 36 hours, limiting glucose availability to T cells (Figure 1h). We validated that the transwell co-culture produced media deficient in glucose available to T cells (Supplemental Figure 1a). Using the Seahorse Bioanalysis Real-Time ATP Rate Assay, we confirmed that T cells co-cultured with tumor cells underwent repressed ATP synthesis that was dependent on glycolysis (glycoATP), and glucose supplementation restored glycoATP in T cells in transwells (Hurst et al., 2020; Riesenberg *et al*., 2022) (Supplemental Figure 1b). These data indicate that the transwell co-culture assay mimics T cells in a fasted state due to low glucose availability that represses T cell energetics. Cells exposed to nutrient stress utilize mitochondrial FAO to sustain ATP synthesis. However, in CD8^+^ T cells exposed to TME-associated nutrient stress we observed that ATP synthesis from the mitochondria (mitoATP) was depressed (Figure 1i). We reasoned that increased expression of ACC1/2 could impede the ability of T cells to utilize FAO-based metabolism. Therefore, we performed western blotting to measure ACC1/2 in T cells harvested from control or tumor-seeded transwells. We observed induction of ACC in T cells co-cultured with tumor cells, relative to expression in tumor-free controls (Figure 1j).

Given that we noted an increase in ACC1/2 in T cells exposed to nutrient stress, we reasoned that overexpression of ACC1/2 could impede the ability of T cells to transition to FAO-based metabolism. Using the ATP rate assay, we measured mitoATP in control and TME-exposed CD8^+^ T cells in the presence or absence of etomoxir, a potent inhibitor of CPT1A that imports free fatty acyl-coA to the inner mitochondrial membrane during the oxidation of fats. We observed that in control conditions CD8^+^ T cells exhibited dependence on FAO for mitoATP synthesis (Figure 1k). In contrast, TME exposure blunted dependence of T cells on FAO for sustained mitoATP (Figure 1l). Steatosis is the accumulation of lipids that culminates in increased lipid storage in the form of lipid droplets (LDs), and can be caused by ACC1/2 activity in the absence of complimentary lipid utilization/ degradation (Kim et al., 2017). The data prompted us to use confocal microscopy to measure LDs in CD8^+^ T cells exposed to nutrient stress. Using the neutral lipid dye Bodipy 493/503, we labeled LDs in relation to the endoplasmic reticulum (ER) (SEC61b), the lipid-rich organelle that generates LDs (Basseri and Austin, 2012), and nucleus (DRAQ5). Indicative of lipid storage, we noted that CD8^+^ T cells exposed to tumor stress harbored increased numbers of LDs on a per cell basis, compared to tumor-free controls (Figure 1m). Next, we probed our *in vivo* RNA-seq data set (Figure 1a) for evidence of lipid storage disorders. We noted that the gene set indicative of LD generation was significantly enriched in CD8^+^ TILs relative to CD8^+^ T cells isolated from spleens (Figure 1n). Thus, we used confocal microscopy to measure LDs in CD8^+^ TILs and autologous splenic CD8^+^ T cells isolated from mice bearing MCA-205 fibrosarcomas. Strikingly, CD8^+^ TILs harbored substantially more LDs relative to splenic T cells (Figure 1o). These data indicate that inhibition of FAO and increased lipid storage correlate with ACC1/2 expression in CD8^+^ T cells exposed to tumor stress.

### ACC inhibition rewires CD8^+^ T cell metabolism

The therapeutic ND-646 was developed as a potent allosteric ACC1/2 inhibitor that binds the biotin carboxylation domain of ACC, thwarting dimerization. Binding of ND-646 leaves ACC serine phosphorylation sites open to constitutive dephosphorylation. Therefore, abrogation of ACC1/2 serine phosphorylation (mouse ACC1 Ser79; human ACC1 Ser117; mouse ACC2 Ser212; human ACC2 Ser222) presents a biomarker to monitor ND-646 target binding efficacy (Svensson *et al*., 2016). We aimed to determine whether ND-646, hereafter referred to as ACCi, could act as a potent inhibitor of ACC in CD8^+^ T cells, as a tool to alter metabolism. We cultured *ex vivo* expanded OT-1 T cells in vehicle or ACCi, then performed immunoblotting for p-ACC using an antibody that recognizes Ser79 phosphorylation. p-ACC signal was abrogated in ACCi-treated T cells, indicating target-specific binding of the drug (Supplemental Figure 2a), and a flow cytometry-based approach to detect p-ACC at Ser79 showed a similar reduction in p-ACC expression in ACCi-treated cells (Supplemental Figure 2b). ACC converts acetyl-CoA to malonyl-CoA, providing 2-carbon units to fatty acids, serving as the first committed step in FAS (Foster, 2012). To determine whether ACC activity was impaired by ACCi, we measured malonyl-coA levels in OT-1 T cells activated and expanded in the presence of vehicle or ACCi. We observed that ACCi substantially reduced malonyl-CoA levels at multiple time points over the course of T cell expansion *in vitro* (Supplemental Figure 2c), indicating consistent limited ACC activity in CD8^+^ T cells. Malonyl-CoA promotes lipogenesis that provides lipids essential for cell membranes, allowing cell growth and division (Lombard, 2014). However, after a 7-day course of OVA-peptide mediated OT-1 T cell activation and expansion in the presence of vehicle or ACCi, we did not observe a significant reduction in total cell numbers between groups (Supplemental Figure 2d), suggesting ACCi did not produce a defect in T cell expansion. Together, these data suggest that ACCi, initially developed to limit FAS in tumor cells (Svensson *et al*., 2016), also limits ACC1/2 activity in CD8^+^ T cells.

Next, we performed metabolomics on peptide-activated OT-1 T cells activated and expanded for 7 days in the presence or absence of ACCi to determine whether the therapeutic could produce metabolic changes in CD8^+^ T cells. Vehicle and ACCi-treated T cell groups exhibited distinct separation by principal component analysis, suggesting that metabolite profiles between groups differed substantially (Figure 2a). Among 213 metabolites detected, 69 were significantly differentially regulated between groups, based on our criteria for significance. Notably, the bulk of significantly differentially regulated metabolites were increased in the ACCi-treated T cell group, as illustrated by volcano plot (Figure 2b). Next, we performed quantitative enrichment analysis (QEA) to identify metabolic pathways enriched in the data set. We observed that mitochondrial β-oxidation of short, medium, and long-chain fatty acids and oxidation of branched chain fatty acids were among the significantly enriched pathways (FDR < 0.05) that comprised the data set (Figure 2c). Enrichment signatures of mitochondrial β-oxidation of short, medium, and long-chain fatty acids as well as oxidation of branched chain fatty acids were driven in part by increased abundance of ATP and acetyl-CoA in ACCi-treated T cells compared to vehicle controls (Figures 2d). Given that acetyl-CoA generated by FAO can enter into TCA, providing NADH and FADH2 equivalents used in turn in the electron transport chain to generate ATP (Martínez-Reyes and Chandel, 2020), we probed the data set for metabolites associated with TCA activity in the T cell groups. Notably, we detected increases in a majority of TCA intermediates in ACCi-conditioned T cells including α-ketoglutarate (α-KG), fumarate, and malate (Figure 2e). Together, the data indicate a substantial metabolic rewiring toward FAO dependence in ACCi-conditioned T cells.

**Figure 2.**
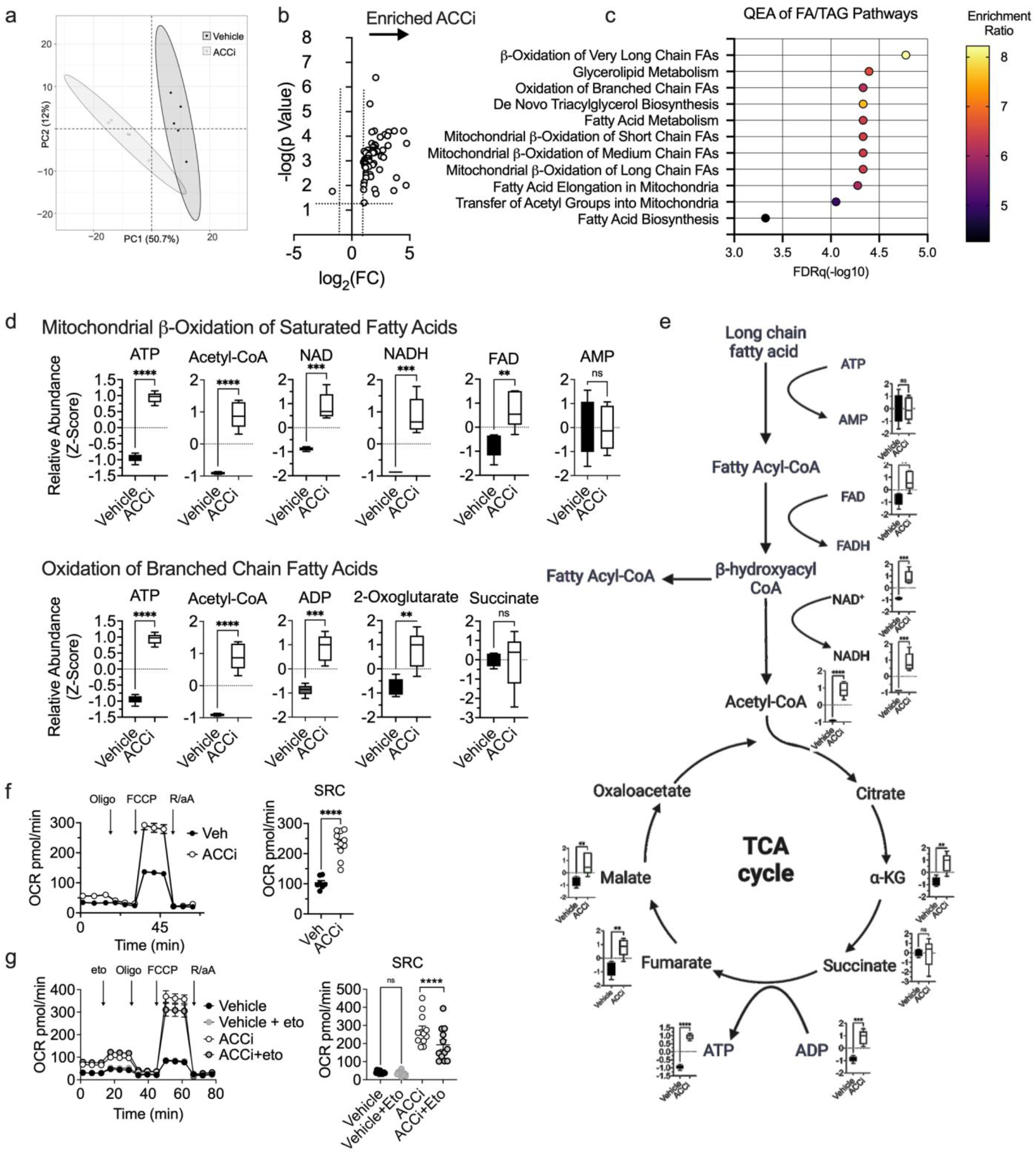
ACC inhibition rewires CD8^+^ T cell metabolism. OT-1 T cells were activated with OVA peptide in the presence of IL-2 and expanded ± ACCi for 7 days. a) Principal component analysis of metabolomics data generated from T cell preparation from 5 mice cultured ± ACCi. b) Volcano plot indicating significantly differentially regulated metabolites between T cell preparations from 5 mice cultured ± ACCi, criteria for significance Adj *p* < 0.05, fold change (FC) > 2. c) Quantitative enrichment analysis (QEA) representing significantly enriched pathways associated with lipid metabolism driven by metabolite levels in T cell preparations from 5 mice cultured ± ACCi. d) Relative abundance of individual metabolites driving QEA signatures (as seen in c) of mitochondrial β-oxidation of short, medium, and long chain fatty acids and oxidation of long chain fatty acids between T cell preparations from 5 mice cultured ± ACCi. e) Schematic of FAO feeding into the TCA with metabolite levels of TCA intermediates identified between T cell preparations from 5 mice cultured ± ACCi. Seahorse Mito Stress Test Assay was performed on OT-1 T cells activated with OVA peptide in the presence of IL-2 and expanded ± ACCi for 7 days. f-g) Oxygen consumption rate (OCR) trace with spare respiratory capacity (SRC) quantification in response to indicated injections of oligomycin (Oligo, ATP synthase inhibitor), FCCP (proton gradient uncoupling), rotenone/antimycin A (R/Aa, electron transfer inhibition) and addition of g, etomoxir (Eto, CPT1A inhibitor). SRC was calculated as the difference between the basal and maximal OCR readings after addition of FCCP. Data points represent individual Seahorse wells. Independent experiments repeated at least 3 times. ns, not significant, * p < 0.05, ** p < 0.01, *** p < 0.001, **** p < 0.0001 based on statistical analysis by unpaired Student’s t-test, all error bars indicate the SEM. Criteria for significance from QEA were FDR Adj p < 0.05, (c).

A hallmark of T cells with FAO dependence is increased spare respiratory capacity (SRC) (van der Windt *et al*., 2012; Windt *et al*., 2013), a parameter measured by Seahorse Bioanalysis that measures reserve capacity of ETC when uncoupled from ATP synthase, indicating an excess of ETC capacity over oxidative phosphorylation. We performed Seashore Mito Stress Test Assay to assess SRC between OT1-T cells activated and expanded for 7 days in the presence or absence of ACCi. We observed that ACCi-conditioned cells harbored increased SRC relative to vehicle controls (Figure 2f). To definitively test whether mitochondrial SRC was dependent on FAO, we performed the Seahorse Mito Stress Test Assay with or without introduction of the CPT1A inhibitor Etomoxir (Eto), to measure SRC in response to inhibition of fatty acyl-coA import to the mitochondria. In the presence of Eto, SRC was not substantially affected in vehicle T cells. However, we observed a significant reduction in SRC in ACCi-conditioned cells treated with Eto, relative to ACCi controls (Figure 2g). These data indicate that the enzyme ACC restricts synthesis of ATP-derived from mitochondrial FAO in CD8^+^ T cells.

### ACC limits energy synthesis in T cells in nutrient stress

How T cells generate energy in the nutrient stress of solid tumors is of great interest to the tumor immunotherapy field in order to more appropriately tune the success of T cell therapies in solid cancers. Given that FAO is a metabolism predominantly employed by cells experiencing nutrient stress, we aimed to study how ACC activity affects ATP synthesis in CD8^+^ T cells exposed to the TME. To measure the effect of ACCi on T cell ATP synthesis in nutrient stress, we harvested T cells from the tumor cell-T cell transwell co-culture assay (Figure 1h) (Hurst *et al*., 2020; Riesenberg *et al*., 2022) and performed Seahorse Bioanalysis Real-Time ATP Rate Assay that measures and discriminates ATP generated within the mitochondria (mitoATP) and ATP synthesized from glycolysis (glycoATP) in cells. We observed that vehicle-treated T cells experienced repression of total ATP synthesis in nutrient stress. In contrast, CD8^+^ T cells conditioned with ACCi exhibited increased total ATP production in nutrient stress, indicating that ACC particularly restricted T cell bioenergetics in a fasted state (Figure 3a). Quantification of mitoATP (Figure 3b) and glycoATP (Figure 3c) suggested that ACC inhibition augmented synthesis of both mitoATP and glycoATP in T cells under nutrient stress.

**Figure 3.**
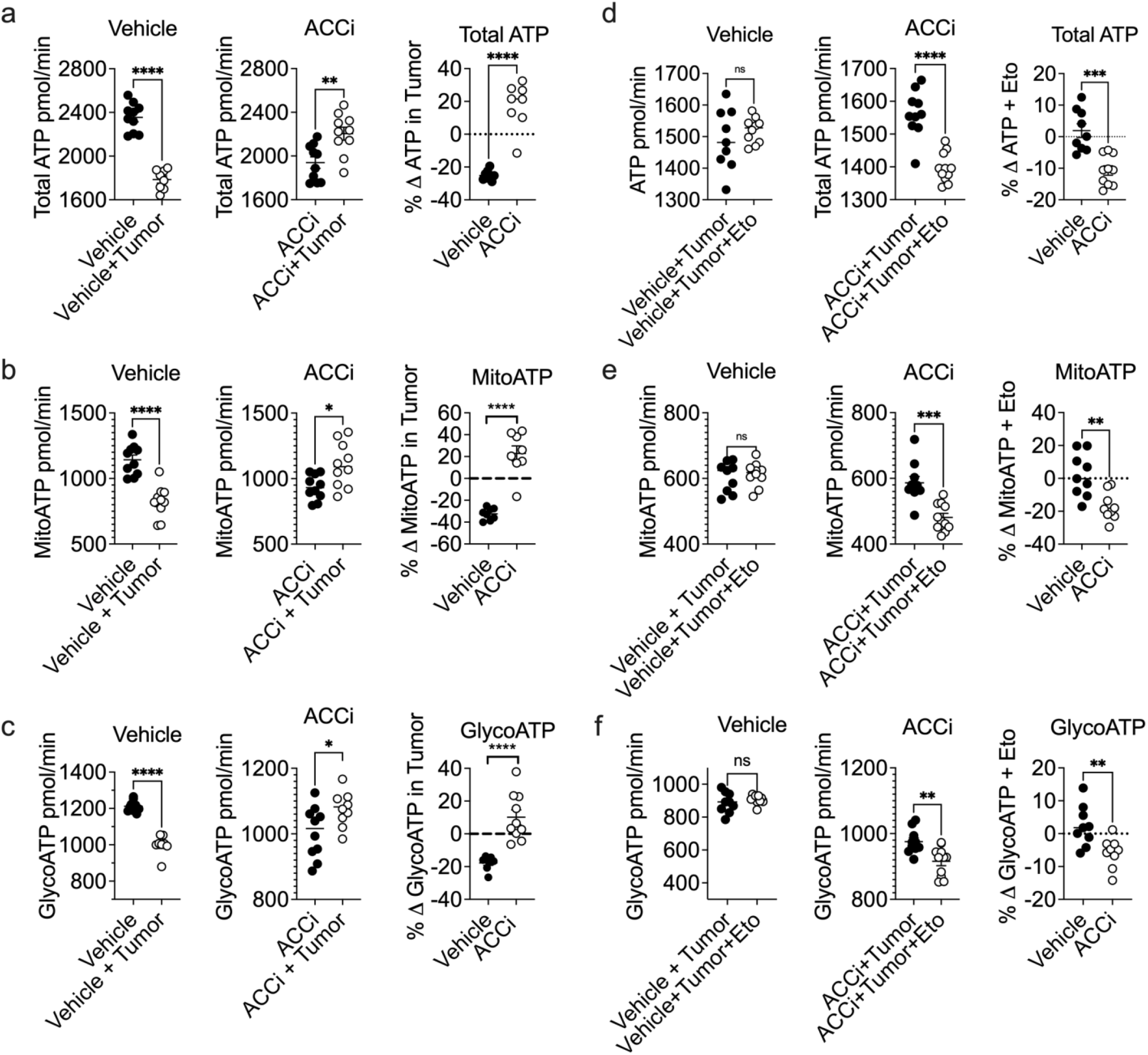
ACC Limits Energy Synthesis in T cells in Nutrient Stress. OT-1 T cells were activated with OVA peptide in the presence of IL-2 and expanded for 3 days then seeded into control or transwells seeded with B16F1 melanoma cells ± ACCi. After 36 hours of co-culture, T cells were collected and Seahorse Real-Time ATP Rate Assay was performed. Quantifications among T cell groups of a,d) total ATP rates b,e) mitochondrial ATP rates (mitoATP) and c,f) glycolytic ATP rates (glycoATP) are shown in response to sequential injections of a-c, oligomycin and rotenone/antimycin A or d-f, etomoxir or buffer control, oligomycin, and rotenone/antimycin A. mitoATP rate was calculated based on the decrease in the OCR. glycoATP rate was calculated as the increase in the ECAR combined with total proton efflux rate. Data points represent individual Seahorse wells. Independent experiments repeated at least 3 times. ns, not significant, * p < 0.05, ** p < 0.01, *** p < 0.001, **** p < 0.0001 based on statistical analysis by unpaired Student’s t-test, all error bars indicate the SEM.

To definitively test whether increased ATP synthesis in ACCi-treated T cells was due to FAO we performed the ATP rate assay with or without introduction of Eto, as previously described (Figure 1). We observed that CPT1A inhibition did not affect total ATP, mitoATP, or glycoATP synthesis in vehicle-treated T cells exposed to nutrient stress. In stark contrast, introduction of Eto significantly repressed total ATP, mitoATP, and glycoATP in ACCi-treated T cells in the fasted state (Figures 3d-f). Taken together, these data indicate that ACC restricts T cell bioenergetics in TME stress, and illustrate that ACC inhibition provides a benefit of improved ATP synthesis to T cells entering the TME environment.

### ACC dictates lipid composition of T cells

The data indicated greater utilization of FFAs in ACCi-conditioned T cells relative to vehicle controls. However, limited ACC activity reduces biosynthesis of lipids (Wakil and Abu-Elheiga, 2009). To rectify these data, we performed global lipidomics analysis to assess lipid content between CD8^+^ T cells expanded in the presence of vehicle or ACCi. We used UPLC-MS-based lipidomics to identify lipid classes in vehicle and ACCi-treated T cells. As expected, we observed a significant reduction in total lipid content measured by integration of total peak area among ACCi-treated T cells relative to vehicle controls (Figure 4a), indicating that limiting ACC activity reduced lipid biosynthesis in T cells. Next, we compared abundance of individual lipid classes between vehicle and ACCi-treated T cells. Among major lipid classes, we observed that triacylglycerides (TAG), diacylglycerides (DAG), phosphatidylcholines (PC), phosphatidylglycerols (PG), and phosphatidylserines (PS) were significantly reduced in ACCi-treated T cells relative to vehicle controls. The changes were indicative of a global deficit in lipid biosynthesis in T cells responding to restricted ACC activity. However, we noted that sphingomyelins (SM) and lysophosphatidylcholines (LPC), a minor lipid class that can be rapidly metabolized (Law et al., 2019), were significantly increased in ACCi-conditioned CD8^+^ T cells compared to vehicle controls (Figure 4b). Together, these data show that limiting ACC promotes abundance of choice lipid species in CD8^+^ T cells.

**Figure 4.**
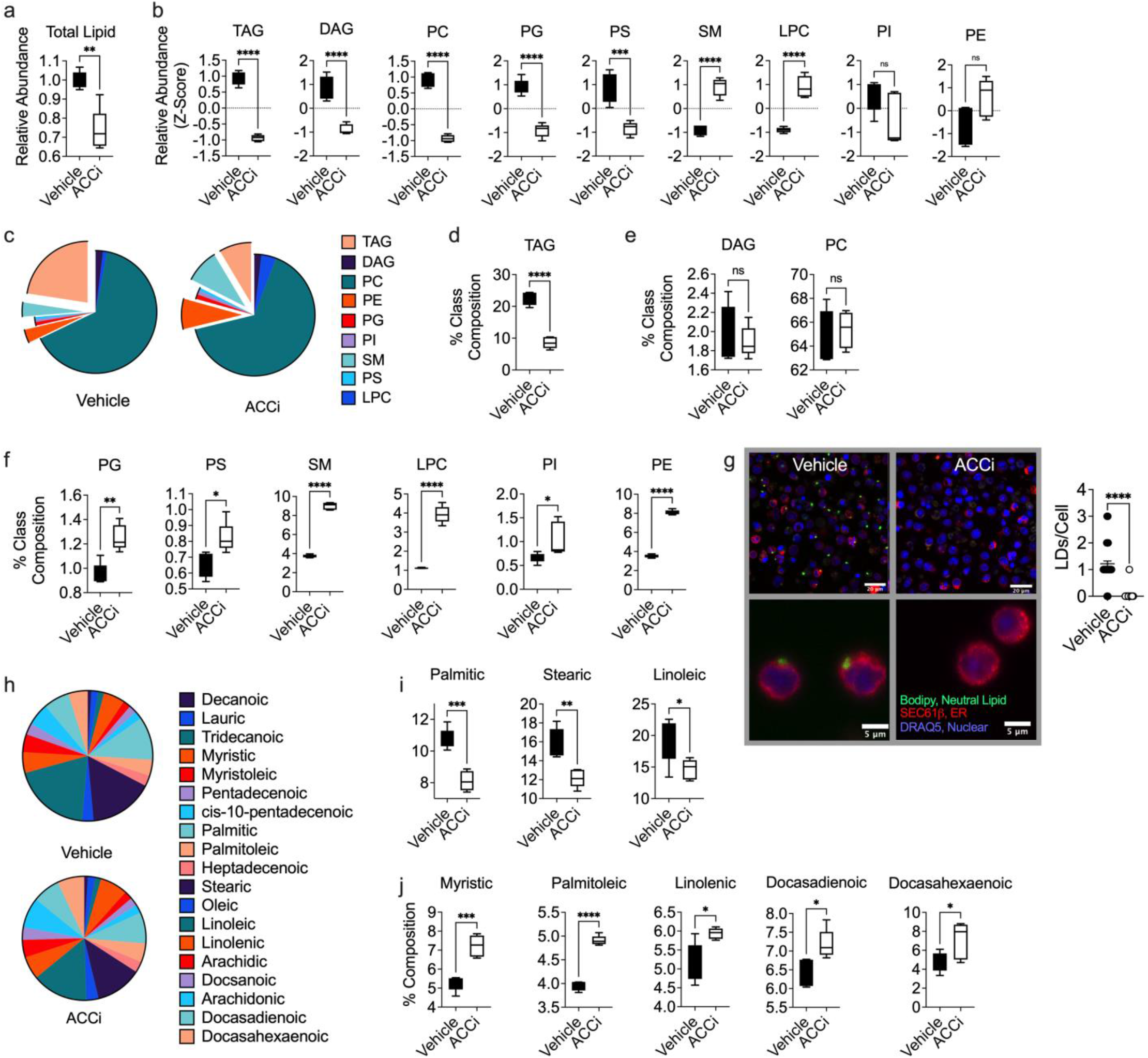
ACC dictates lipid composition of T cells. OT-1 T cells were activated with OVA peptide in the presence of IL-2 and expanded ± ACCi for 7 days. UPLC-MS lipidomics was performed on T cell cultures from n=5 mice treated ± ACCi. a) Quantification of integration of total peak area of lipids detected by UPLC-MS. b) Relative abundance of individual lipid classes identified between vehicle and ACCi-treated T cell groups. c) Pie chart representations of average percent lipid class composition between vehicle or ACCi-treated T cell groups. Quantification of d) significantly decreased (TAG) e) not significantly regulated (DAG, PC) or f) significantly increased (PG, PS, SM, LPC, PI, PE) major lipid classes measured by percent lipid class composition between vehicle or ACCi-treated T cell groups. Cells were probed with Bodipy 493/503 and confocal microscopy was used to g) visualize and quantify stored neutral lipids in the form of LDs between vehicle or ACCi-treated T cell groups. g, Data points represent number of lipid LDs per cell (n=50). GC-MS was used to perform targeted FFA analysis on T cell cultures from n=5 mice treated ± ACCi. h) Pie chart representations of average percent FFA composition between vehicle or ACCi-treated T cell groups. Significantly increased FFAs identified in i) vehicle and j) ACCi-treated T cell groups as measured by percent composition. Independent experiments repeated at least 2 times. FFA analysis performed once. ns, not significant, * p < 0.05, ** p < 0.01, *** p < 0.001, **** p < 0.0001 based on statistical analysis by unpaired Student’s t-test, all error bars indicate the SEM.

Though the data suggested that the majority of lipid classes were diminished by ACC inhibition, we questioned whether the percent composition of each lipid class comprising each T cell group was similar. We first visualized each lipid class as a percentage of total lipids contained in vehicle or ACCi-treated T cells using pie charts. We observed that PCs were the most abundant lipid class in CD8^+^ T cells, comprising ∼60% of total lipids, and TAGs were the second most abundant lipid class covering ∼20% of the lipidome. Pie chart visualization indicated that, relative to vehicle, ACCi-conditioned T cells harbored fewer TAGs, but contained more SMs and PEs (Figure 4c). Indeed, quantification of percent lipid class composition identified that TAGs were the only lipid class significantly reduced by ACC inhibition (Figure 4d). While DAGs and PCs comprised a similar proportion of the total lipidome in both cell groups (Figure 4e), the lipidome of ACCi-treated T cells harbored greater proportions of PG, PS, SM, LPC, PI and PE lipid classes (Figure 4f). These data establish that ACC activity dictates lipid composition in CD8^+^ T cells.

In nutrient replete states, TAGs are stored in cytoplasmic LDs (Rambold et al., 2015). Such neutral lipid stores can be detected and quantified by flow cytometry or confocal microscopy to assess neutral lipids on a per cell basis or to visualize the spatial location of LDs, respectively. Given that lipidomics showed that TAGs were the main lipid class in reduced proportions relative to vehicle controls, we aimed to assess TAG content between the two T cell groups. We labeled CD8^+^ T cells with Bodipy 493/503, DRAQ5 nuclear stain, or SEC61β to locate the ER relative to TAGs. Strikingly, clear LDs were visible in vehicle control cells, but LDs were notably absent from ACCi-treated T cells (Figures 4g), suggesting that ACC restriction resulted in loss of lipid storage in CD8^+^ T cells.

FFA pools can be stored in LDs. Given that vehicle T cells harbored increased LD stores, we reasoned that ACC inhibition could promote an increased pool of FFAs available for mitochondrial consumption as a potential mechanism for increased FAO in ACCi T cells. We used gas chromatography-liquid mass spectrometry (GC-MS) to perform targeted FFA analysis between the T cell groups, but did not observe significant differences in total amount of the FFA pool between T cell groups. Next, we projected the 19 types of FFAs identified as percent composition of each T cell group onto pie charts (Figure 4h) to visualize whether one T cell pool was enriched in a specific FFA type. Percent composition analysis indicated that vehicle T cells were enriched for palmitic, stearic, and linoleic acids (Figure 4i), indicating that vehicle T cells were enriched in saturated FFAs (palmitic, stearic) compared to ACCi T cells. These data corroborated the finding that vehicle T cells harbored increased malonyl-coA (Supplemental Figure 2), as malonyl-coA provides the substrate for palmitate. In contrast, ACC inhibitor-treated T cells were enriched in 5 FFA types, four of which constituted unsaturated FFAs, including palmitoleic acid, docasadienoic acid, linolenic acid, and docasahexaenoic acid (DHA), purported to be rapidly metabolized to promote oxidative phosphorylation and ATP synthesis (Cruz et al., 2018; Ide et al., 1996). These data suggest that ACC competency enforces lipid storage in T cells, obstructing fat utilization.

### ACC obstructs ER-Directed FFA utilization

The data prompted us to question the cellular resource of FFAs in CD8^+^ T cells responding to ACC restriction. Though ACCi-conditioned T cells harbored fewer lipids, and were comprised of a reduced proportion of TAGs, we noted that *de novo* TAG biosynthesis was significantly enriched in the metabolomic data set (Figure 2). Further examination indicated that metabolites driving *de novo* TAG biosynthesis were significantly increased in the ACCi-conditioned T cell group relative to vehicle controls (Supplemental Figure 3). In the ER, enzymes that comprise the glycerol 3-phosphate (G-3-P) pathway carry out *de novo* TAG biosynthesis, leading to subsequent TAG storage in LDs (Harris et al., 2011). In fasted states LDs are liberated by lipolysis to access FFAs that can fuel mitochondrial energetics (Rambold *et al*., 2015). Thus, ER directed TAG biosynthesis presents a possible resource of FFAs. The enzyme diglyceride acyltransferase (DGAT) catalyzes the committed step of TAG synthesis in the ER (Figure 5a) (Harris *et al*., 2011). Metabolomics analysis suggested that glycerolipid metabolism was enriched in the data set (Figure 1), and the metabolites driving the enrichment signature were more abundant in ACCi-treated T cells. Among such metabolites, NADH and dihydroxyacetone phosphate (DHAP) provide the substrates to generate G-3-P, and increased glyceric acid, the oxidized form of glycerol, were all elevated in ACCi-conditioned T cells relative to vehicle controls (Figure 5b). Next, we performed RNA-sequencing to determine whether the differential metabolism in response to ACCi-conditioning was associated with genetic reprogramming of glycerolipid synthesis in the CD8^+^ T cells. We observed that genes encoding enzymes required for glycerolipid metabolism and subsequent TAG synthesis in the ER (Figure 5a) were profoundly upregulated in ACCi-conditioned T cells relative to vehicle controls (Figure 5c).

**Figure 5.**
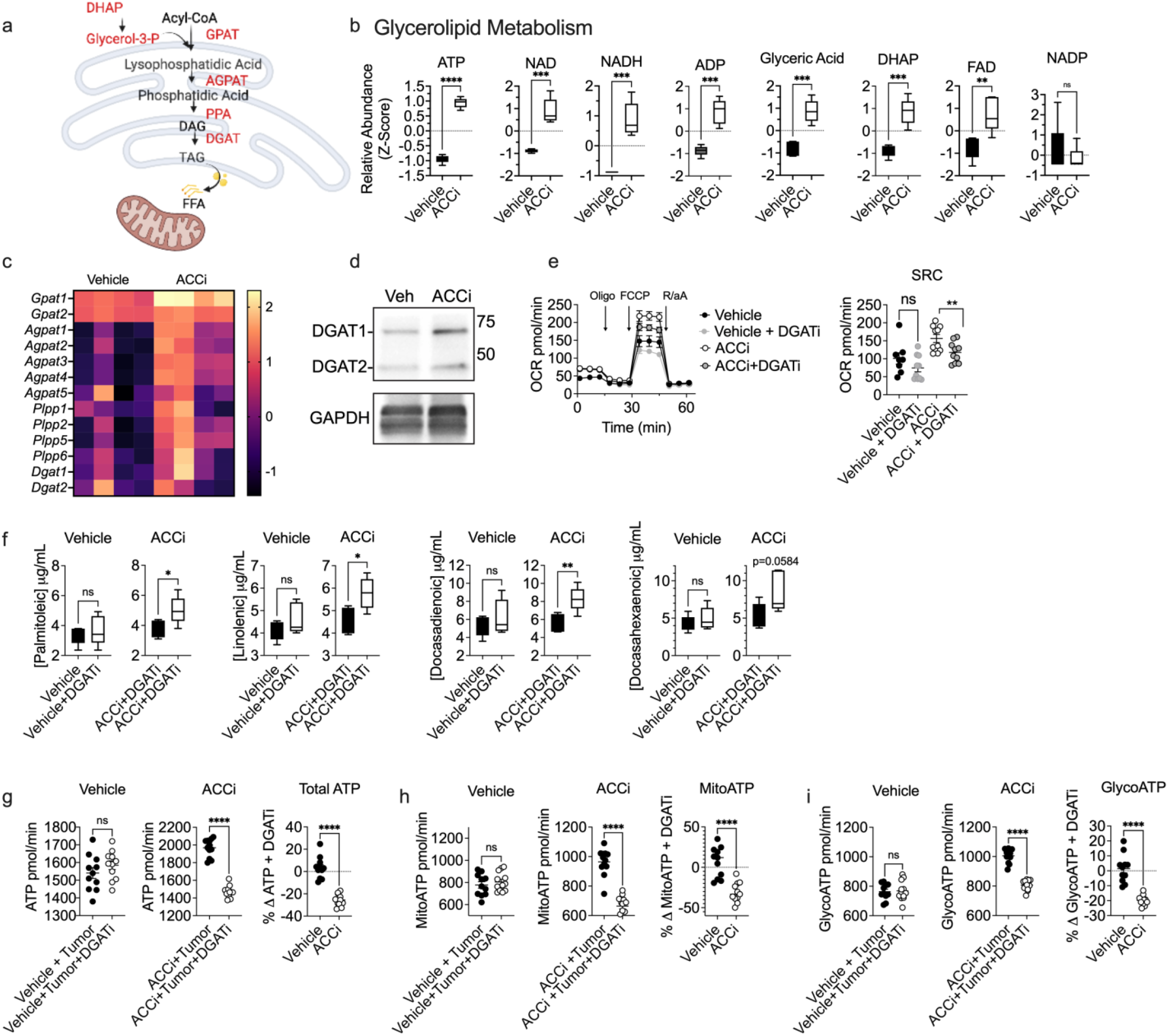
ACC obstructs ER-Directed FFA utilization. OT-1 T cells were activated with OVA peptide in the presence of IL-2 and expanded ± ACCi for 7 days. a) Schematic of glycerol-3-phosphate (G-3-P) pathway in the ER that generates triacylglycerides (TAGs) via the enzyme diglyceride acyltransferase (DGAT). The pathway can serve as a source of FFAs. b) Relative abundance of individual metabolites driving QEA signature for Glycerolipid Metabolism between T cell preparations from 5 mice cultured ± ACCi, analysis and significance as described in Figure 1. c) Heat map of normalized reads from RNA-sequencing of genes encoding isoforms of the major metabolic enzymes in the G-3-P pathway as depicted in a, between T cell preparations from 4 mice cultured ± ACCi. d) Western blotting for DGAT enzyme that catalyzes the committed step in TAG synthesis in the ER as depicted in a, in T cells cultured ± ACCi. OT-1 T cells were activated with OVA peptide in the presence of IL-2 and expanded ± ACCi for 7 days ± DGATi. e) Seahorse Mito Stress Test Assay was performed to measure oxygen consumption rate (OCR) trace with spare respiratory capacity (SRC) quantification in response to indicated injections of oligomycin (Oligo, ATP synthase inhibition), FCCP (proton gradient uncoupling), rotenone/antimycin A (R/Aa, electron transfer inhibition). SRC was calculated as the difference between the basal and maximal OCR readings after addition of FCCP. Data points represent individual Seahorse wells. FFA analysis was performed and f) quantification of FFAs identified as significantly enriched in ACCi-treated T cells in Figure 4j is shown. OT-1 T cells were activated with OVA peptide in the presence of IL-2 and expanded for 3 days then seeded into control or transwells seeded with B16F1 melanoma cells ± ACCi ± DGATi. After 36 hours of co-culture, T cells were collected and Seahorse Real-Time ATP Rate Assay was performed. Quantifications among T cell groups of f) total ATP g) mitochondrial ATP (mitoATP) and h) glycolytic ATP rates (glycoATP) are shown in response to sequential injections of oligomycin and rotenone/antimycin A. mitoATP rate was calculated based on the decrease in the OCR. glycoATP rate was calculated as the increase in the ECAR combined with total proton efflux rate. Data points represent individual Seahorse wells. Independent experiments repeated at least 3 times. FFA analysis performed once. ns, not significant, * p < 0.05, ** p < 0.01, *** p < 0.001, **** p < 0.0001 based on statistical analysis by unpaired Student’s t-test, all error bars indicate the SEM.

To determine whether the DGAT-directed TAG synthesis pathway could have functional implications in T cells responding to ACC restriction, we used western blotting to probe for DGAT expression between vehicle and ACCi-conditioned T cells. In line with RNA-seq gene expression data (Figure 5c), we observed that the DGAT1 isoenzyme was increased in ACCi-conditioned T cells relative to vehicle controls (Figure 5d), suggesting increased utilization of the TAG synthesis pathway. To test whether *de novo* TAG synthesis provided a resource of FFAs to CD8^+^ T cells in response to ACC blockade, we performed the Seahorse Mito Stress Test Assay on vehicle and ACCi-conditioned T cells in the presence or absence of Amidepsine A (DGATi), a potent inhibitor of TAG formation through blockade of DGAT enzymatic activity (Tomoda et al., 1995). We found that DGATi significantly reduced reserve mitochondrial ATP synthesis in ACCi-conditioned T cells, but vehicle-treated T cells were unaffected (Figure 5e). The data prompted us to test whether DGAT activity in the ER acted to supply the FFAs that were significantly enriched in ACCi-conditioned cells (Figure 4) for mitochondrial FAO. Thus, we performed targeted FFA analysis on vehicle and ACCi-conditioned T cells treated with vehicle or DGATi. Strikingly, DGATi promoted accumulation of each of the four significantly upregulated unsaturated FFAs in ACCi-treated T cells, but vehicle T cells were unaffected by DGAT inhibition (Figure 5f). Together, these data suggest that DGAT enzymatic activity is required to enable robust mitochondrial utilization of unsaturated FFAs for FAO, generated in response to ACC restriction.

Our prior data suggested that CD8^+^ T cells accelerate utilization of FFAs for fuel in TME nutrient stress (Figure 3), as ATP rates were increased in ACCi-treated T cells in the TME relative to unstressed ACCi controls. Moreover, we found that the effect was dependent on FAO, as inhibition of CPT1A abrogated the effect. Thus, we tested whether the effect was dependent on ER TAG synthesis as the resource that supplied FFAs to mitochondria in nutrient stress. As in Figure 3, we performed the Seahorse Real-Time ATP Rate Assay on T cells harvested from the tumor cell-T cell transwell co-culture assay conditioned ± ACCi ± DGATi. Compared to ACCi control cells, we observed that ACCi cells treated with DGATi displayed a substantial deficit in total ATP, mitoATP, and glycoATP synthesis (Figure 5g-i), but limiting DGAT in vehicle T cells did not substantially affect ATP rates. Together, these data suggest that ACC restriction forces reliance on ER-directed TAG synthesis to support bioenergetics.

### ACCi Promotes T Cell Immunity Against Tumors

A prior report that indicated DGAT expression promoted human memory T cell development (Cui et al., 2015), prompted us to study the effect of ACCi conditioning on T cell lineage traits. Gene Set Variation Analysis performed on RNA-sequencing data sets (Figure 5) showed that CD8^+^ T cells activated and expanded in the presence of ACCi displayed significant enrichment of genes sets associated with T cell memory traits and diminished signatures of T cell exhaustion relative to vehicle controls (Figure 6a). Next, we used a spectral flow cytometry panel to profile the expression of 18 phenotypic markers on CD8^+^ T cells activated and expanded in the presence of vehicle or ACCi. Dimension reduction analysis using the Uniform Manifold Approximation and Projection (UMAP) algorithm illustrated substantial phenotypic remodeling between vehicle and ACCi treatment groups (Figure 6b). We next projected mean fluorescent intensities (MFIs) of individual parameters onto the UMAP space to discern which markers were responsible for driving the differential phenotypes. We observed that CD8^+^ T cells activated and expanded in the presence of ACCi displayed substantial phenotypic remodeling that encompassed significant upregulation of stemness markers such as CD62L, Tcf1, and Bcl2 relative to vehicle controls (Figure 6c, Supplemental Figure 4).

**Figure 6.**
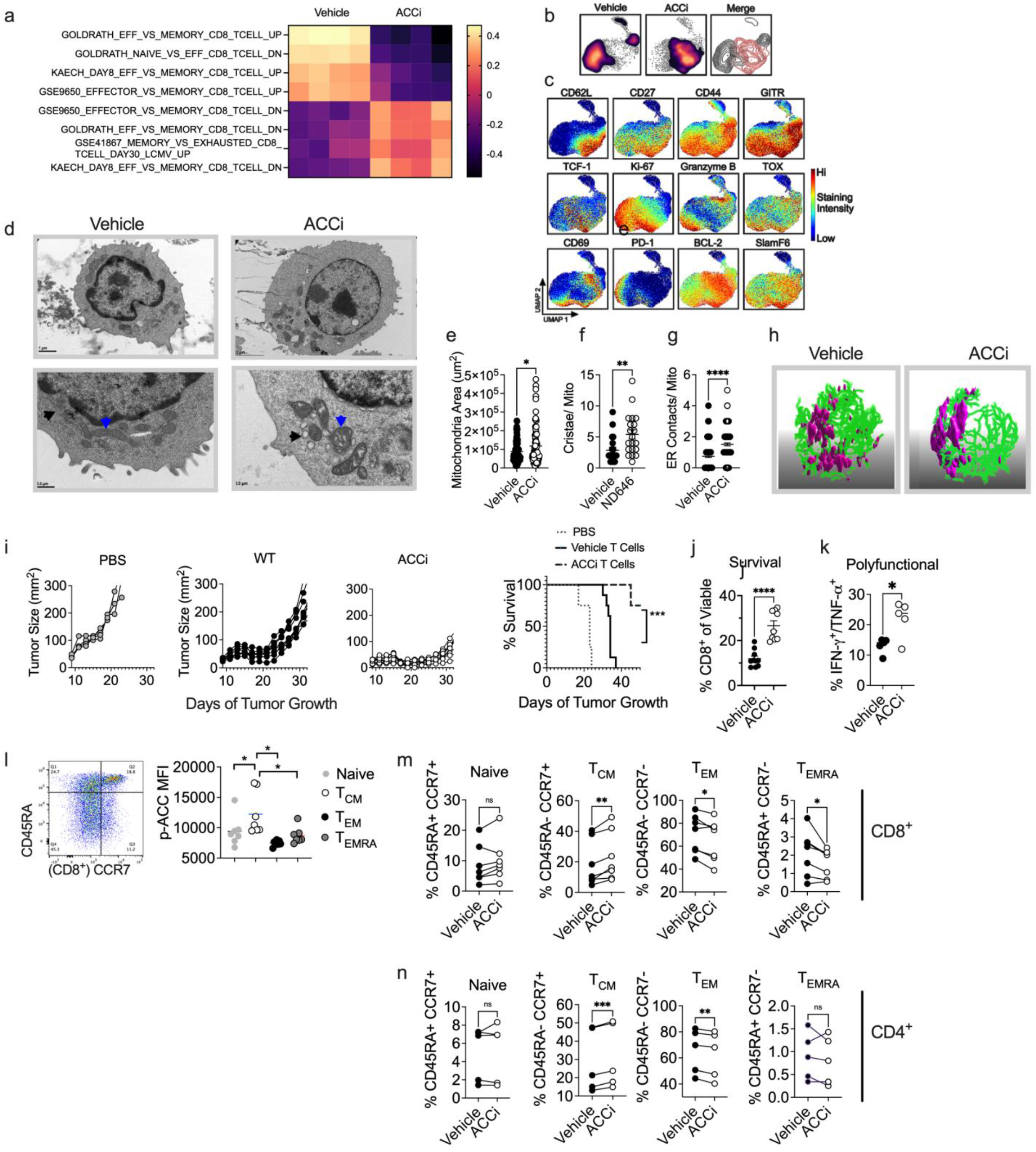
ACCi promotes T cell immunity against tumors. OT-1 T cells were activated with OVA peptide in the presence of IL-2 and expanded ± ACCi for 7 days. a) Heat map of significantly regulated gene sets associated with T cell effector or memory lineages or T cell exhaustion, generated from RNA-sequencing of samples as described in Figure 5. Spectral flow cytometry was performed to assess 18-parameters descriptive of T cell activation, differentiation, and exhaustion and b) dimension reduction was performed using the UMAP algorithm to visualize differences in T cell phenotypes between T cell groups followed by c) projection of mean fluorescent intensities (MFIs) of individual markers onto the UMAP space. d) Representative transmission electron microscopy (TEM) images of vehicle and ACCi-treated T cells. Blue arrows indicate mitochondria, black arrows indicate ER. Quantification of TEM to measure e) mitochondrial area f) cristae per mitochondria g) ER contacts per mitochondria. e-f, Data points represent quantification of individual mitochondria, g, data points represent number of MERCs per cell, n=50 cells. h) Representative 3-D models generated using Imaris software from Z-stacks of confocal imaging performed to visualize mitochondria (purple, TOM20) and ER (green, SEC61β) in vehicle or ACCi-treated T cells. CD45.2^+^ OT-1 T cells activated with OVA peptide in the presence of IL-2 and expanded ± ACCi for 7 days were infused into CD45.1^+^ mice bearing B16F1-OVA melanomas and i) tumor growth and survival were monitored. In a separate group of animals, tumors were harvested 7-days post transfer and j) percentage of viable CD45.2^+^ CD8^+^ T cells and k) polyfunctional cytokine synthesis between transferred T cell groups were quantified. Data points represent averages of n=4-8 mice per group. Independent experiments repeated at least twice. * p < 0.05, ** p < 0.01, *** p < 0.001, **** p < 0.0001 based on statistical analysis by i, log-rank, mantel-cox test of survival proportions and e-g, j-k, unpaired Student’s t-test, all error bars indicate the SEM. For GSVA, criteria for signature enrichment were FDR < 0.05. l) Representative FACS plot and quantification of p-ACC expression among human CD8^+^ T cell subsets. Data points represent individual normal donors (n=7). PBMCs were expanded ± ACCi for 6 days then percentages of m) CD8^+^ or n) CD4^+^ T cell subsets within the cultures were assessed. Naïve, CD45RA^+^/CCR7^+^, T_CM_, CD45RA^-^/CCR7^+^, T_EMRA_, CD45RA^+^/CCR7^-^, T_EM_, CD45RA^-^/CCR7^-^. Data points represent individual human donors. Independent experiments repeated twice. ns, not significant, * p < 0.05, ** p < 0.01, *** p < 0.001 based on statistical analysis by paired Student’s t-test, all error bars indicate the SEM.

Bcl2, found at mitochondria-ER contact sites (MERCs), potentiates maintenance of mitochondria structural dynamics (Aouacheria et al., 2017; Hunt et al., 2022). Given that ACCi resulted in differential ER-mitochondria crosstalk evidenced by increased utilization of ER-derived lipid (Figure 5), we performed transmission electron microscopy (TEM) to visualize organelle structure between vehicle and ACCi T cell groups. We observed differences in mitochondrial morphology in T cells restricted in ACC activity. ACCi-conditioned T cells harbored larger mitochondria (Figures 5d-e), more cristae per mitochondria (Figures 6d, 6f), and increased MERCs (Figures 6d, 6g), suggesting that ACC inhibition prompted mitochondrial health and ER-mitochondrial crosstalk. Notably, in ACCi-conditioned T cells, we observed extensive ER curvature, indicative of lipid producing smooth ER (Spacek and Harris, 1997) (Figure 6d). This differential ER morphology could be necessary for increased TAG synthesis in ACCi-conditioned T cells. To visualize the organellar remodeling induced by ACC limitation in CD8^+^ T cells, we used Z-stacks of confocal imaging coupled with Imaris software to generate 3-D models of vehicle and ACCi T cells. 3-D imaging supported our TEM and western blotting findings, suggesting that ACCi-conditioned T cells harbored larger mitochondria (purple, TOM20) and increased ER curvature (green, SEC61β), compared to vehicle controls (Figure 6h). Together, these images suggest that ACC activity in T cells alters organelle morphology.

T cells with traits of stemness exhibit improved and durable tumor control relative to effector counterparts (Charo et al., 2005; Klebanoff *et al*., 2005; Riesenberg *et al*., 2022). We aimed to test whether T cells conditioned with ACCi supported superior tumor control relative to vehicle controls. We infused OT-1 CD8^+^ T cells activated and expanded in the presence or absence of ACCi into mice bearing B16F1-OVA melanomas and measured tumor control and survival over time. We observed that ACC inhibition improved T cell tumor control and significantly extended time to sacrifice in tumor-bearing hosts (Figure 6i). Next, we infused vehicle or ACCi-treated OT-1 T cells into mice bearing B16F1-OVA melanomas and assessed viability and phenotypic traits of transferred TILs harvested from tumors 7 days after infusion. We observed that T cells rewired by ACCi showed ∼3-fold increase in percentage of viable CD8^+^ TILs accumulating in tumors (Figure 6j) and supported increased polyfunctional cytokine synthesis (Figure 6k).

Our *in vivo* findings prompted us to assess regulation of ACC in human T cells to determine whether ACC inhibition could similarly endow human T cell subsets with differential phenotypic traits, as a possible therapeutic avenue to rewire T cell products for cancer patients. First, we measured the basal level of p-ACC as a mark of ACC restriction in human T cell subsets. We performed intracellular phospho-flow cytometry paired with subset analysis of CD8^+^ T cell populations from peripheral blood mononuclear cells (PBMC) from 7 healthy human donors. Consistent with our findings in murine T cells, among the samples assessed, effector memory RA^+^ (T_EMRA_) and effector memory (T_EM_) CD8^+^ T cell populations had reduced p-ACC expression; however, naïve and central memory (T_CM_) CD8^+^ T cell populations showed increased expression of p-ACC, and p-ACC levels in T_CM_ cells were significantly increased compared to all subsets assessed (Figure 6l). Next, we aimed to test whether ACCi was a tool to promote memory traits in human CD8^+^ T cells. We activated and expanded PBMCs from 5 healthy human donors with soluble anti-human CD3 and anti-human CD28 monospecific antibody complexes then cultured the expanded T cell pool in vehicle or ACCi and used flow cytometry to probe the phenotype of conditioned T cells. In human CD8^+^ T cells, ACCi significantly reduced proportions of T_EMRA_ and T_EM,_ and promoted T_CM_ populations (Figure 6m). Notably, ACC1 restricts formation of T cell memory in human CD4^+^ subsets (Endo *et al*., 2019), suggesting the possibility that ACCi could rewire both CD8^+^ and CD4^+^ T cell populations. In CD4^+^ T cell populations we observed that the proportion of T_EM_ was reduced by ACCi treatment, while the T_CM_ proportion was significantly increased (Figure 6n). Collectively, the data indicate that metabolic rewiring through ACC inhibition is a potential strategy to alter human immunity that could improve outcomes of cellular therapies for cancer patients.

## Discussion

In nutrient stress, the energy sensor AMP-activated protein kinase (AMPK) sustains cell survival and function through inhibitory phosphorylation of ACC (p-ACC), prompting a reduction in FAS, prompting lipid utilization in the mitochondria (Jeon et al., 2012). Activation of AMPK is mediated by a rise in AMP/ATP ratios concomitant with energy stress in the cell, enforcing p-ACC which restricts FAS, sending the cell into a state of lipid catabolism in order to maintain energy synthesis in starvation (Hardie, 2003). Though the TME is state of energy stress, multiple reports show that TILs are unable to sustain ETC activity and ATP synthesis in solid cancers (Chamoto *et al*., 2017; Scharping *et al*., 2016). Though energy synthesis in TILs is repressed, we have previously verified that direct activation of AMPK promotes enhanced T cell function against tumors. Also, we previously noted that exposure to the TME causes robust downregulation of p-AMPK in T cells, serving as a potential mechanism of repressed ATP synthesis of T cells in tumors (Hurst *et al*., 2020). However, the specific contribution of ACC was not studied. Here, using murine and human samples we identify that ACC1 is robustly induced in TILs relative to expression in autologous splenocytes or peripheral T cells, respectively (Figure 1). These data fill in the gap in knowledge regarding repression of ATP synthesis in CD8^+^ TILs, as the paradoxical upregulation of ACC in nutrient stress provides a direct mechanism, not only lack of FAO utilization in TILs, but for staunch repression of FAO for energy synthesis in TILs.

Unexpectedly, we observed that T cells exposed to the TME produce more ATP when ACC activity is restricted than T cells in control conditions (Figure 3). These data further support the concept that limiting ACC is vital to accessing ATP synthesis once in the solid TME, likely due to incapacity to rely on import of glucose for glycoATP once in nutrient stress. We made the curious observation that T cells restricted in ACC in the TME not only enhance mitoATP, but increase glycoATP synthesis as well, both above ACCi T cells in control conditions as well as above vehicle upon TME exposure. A plausible explanation for this is that ACC inhibition promotes glycogen stores through gluconeogenesis that enable acceleration of glycogenolysis in tumor stress. We noted increased glycogen in ACCi-treated T cells (data not shown), a trait previously correlated to T cell memory and enhanced longevity *in vivo* (Ma et al., 2018). From a mechanistic standpoint, FAO produces mitochondrial acetyl-CoA used to turn the TCA, enabling generation of malate which can be metabolized in the cytosol to oxaloacetate and subsequently phosphoenolpyruvate, supporting gluconeogenesis (Zhang et al., 2018). In line with this mechanism, we observed significantly upregulation of both acetyl-CoA and malate in ACC inhibitor-treated T cells. Thus, glycogen stores could provide a resource of intracellular glucose for utilization in nutrient stress.

Though it is established that FAO-enriched T cells promote cancer control, little attention has been paid to the type, quality, and source of FFAs utilized in the process. Recently, treatment of CAR-T cells with polyunsaturated FFA linoleic acid promoted tumor control (Nava Lauson et al., 2023). Here, FFA analysis indicated that ACC restricted the enrichment of monounsaturated FFAs in T cells, indicating that ACC activity specifically promoted accumulation of saturated FFAs which readily form solid lipid deposits, limiting the ability to be rapidly metabolized (van Marken Lichtenbelt et al., 1997). Specifically, the saturated FFAs palmitic acid and stearic acid were two of the three enriched FFAs in ACC-competent T cells (Figure 4). In stark contrast, limiting ACC prompted T cell access to “healthy fats” able to be readily metabolized for energy including monounsaturated palmitoleic acid, di-unsaturated docasadienoic acid, polyusaturated linolenic acid, and DHA, the highly unsaturated omega-3-fatty acid. These data raise the underexplored possibility that tuning the FFA pool in TILs dicates T cell efficacy against tumors. The FFA pool could be dictated both intracellular regulation of ACC, as well as by the extracellular tumor milieu.

Prior reports in the T cell field determined that memory T cells utilize TAG synthesis to support FAO, and identified that DGAT was enriched in such T cell pools (Cui *et al*., 2015; O’Sullivan et al., 2014). However, these studies were performed in nutrient replete media and did not address T cell ATP synthesis in nutrient stress, or the TME, nor was ACC identified as the upstream metabolic switch that programs TAG store as opposed to utilization. Here, we noted that inhibition of DGAT caused accumulation of unsaturated fats in ACCi T cells, indicating that the committed step of TAG synthesis in the ER was required for the mitochondria to access these fats. This was evidenced by a reduced in FAO in response to DGAT inhibition in the absence of ACC activity. These data raise the questions; How do ACCi T cells produce highly unsaturated fats? As well as, why are FFAs not stored in the context of limited ACC activity? Answers to these questions to elucidate the role of lipid metabolism in antitumor immunity.

Finally, the organellar structural differences between vehicle and ACCi-conditioned T cells shed light on the importance ER-mitochondrial contacts previously identified to be enriched in memory T cells (Hunt *et al*., 2022). It is striking that modulation of the activity of a singular enzyme can reshape both the complete cellular lipidome and organelle structure in T cells. While mitochondrial morphology in T cells has been studied in-depth, ER structure remains largely untouched. Between T cell groups we noted considerable differences in ER morphology, with ACCi-conditioned T cells exhibiting extensive ER curvature that is indicative of lipid-producing smooth ER (Spacek and Harris, 1997). These data compliment the finding that ACC inhibition promoted TAG synthesis in the ER. These data indicate that the interplay between immune cell metabolism, organelle structure, and immune lineage and function is an area to address as a future possibly to reshape the efficacy of clinical immunotherapies.

## Materials & Methods

*Lead contact:* Further information and requests for resources and reagents should be directed to the lead contact Jessica Thaxton.

*Materials availability:* This study did not generate new unique reagents.

*Data and code availability:* All data reported in this paper will be shared by the lead contact upon request.

### Mice

OT-1 (C57BL/6-Tg(TcraTcrb)1100Mjb/J), CD45.1 (B6.SJL-*Ptprc^a^Pepc^b^*/BoyJ), and C57BL/6 mice were obtained from the Jackson Laboratories. All animal experiments were performed in accordance to protocol 21-228 approved by the Institutional Animal Care and Use Committee (IACUC) at the University of North Carolina at Chapel Hill-CH. Mice were housed in a pathogen-free animal facility and maintained by the Division of Comparative Medicine at UNC-CH. Age-matched (6-8 weeks) female mice were used in all mouse experiments. The number of animals (biological replicates) are indicated in figure legends.

### Human Samples

Patients undergoing surgical removal of high-grade deep pleomorphic undifferentiated sarcoma granted informed written consent under the Medical University of South Carolina Biorepository Shared Resource surgical consent forms. This work was determined by the Medical University of South Carolina Institutional Review Board to be exempt under protocol Pro00055960. Tissue and blood samples were deidentified and therefore the sex or gender of participants is unknown. Studies were conducted in accordance with the Declaration of Helsinki, International Ethical Guidelines for Biomedical Research Involving Human Subjects (CIOMS), Belmont Report, or U.S. Common Rule. Blood (4-8 mL) was collected in EDTA coated tubes and peripheral blood mononuclear cells (PBMCs) were isolated via Histopaque-1077 centrifugation (Sigma). Sarcoma tissue was collected on ice and immediately cut into 2 mm^3^ segments and dissociated to a single-cell suspension using the Human Tumor Dissociation Kit and gentleMACS dissociator (Miltenyi Biotec) according to the manufacturer’s protocol. Cells were enriched for CD8^+^ population using the EasySep Human CD8^+^ T Cell Isolation Kit (Stemcell Technologies) and CD8^+^ cells were used for analysis.

### Cell Cultures

MCA-205 (Sigma), B16F1 (ATCC), and B16F1-OVA (kind gift of Dr. Mark Rubinstein) tumor lines and OT-1 T cells were maintained in RPMI supplemented with 10% FBS, 300 mg/L L-glutamine, 100 units/mL penicillin, 100 µg/mL streptomycin, 1mM sodium pyruvate, 100µM NEAA, 1mM HEPES, 55µM 2-mercaptoethanol, and 0.2% Plasmocin mycoplasma prophylactic (InvivoGen). 0.8 mg/mL Geneticin selective antibiotic (Invitrogen) was added to media of B16F1-OVA cells for multiple passages then cells were passaged once in Geneticin-free media prior to tumor implantation. MCA-205, B16F1, and B16F1-OVA cell lines were determined to be mycoplasma-free in March of 2023. For OT-1 T cell activation and expansion, whole splenocytes from OT-1 mice were activated with 1 µg/mL OVA 257-264 peptide (InvivoGen) and expanded for 3 days with 200 U/mL rhIL-2 (NCI) ± 100nM ACCi (ND-646, MedChemExpress), and in some cases ±20μM DGATi (Amidepsine A, Cayman Chemical) then split and expanded into similar conditions for 4 more days. In some experiments, 3-day expanded OT1 T cells were treated ± ACCi and harvested after 36 hours of coculture in transwell assays. Human PBMCs were activated with Immunocult^TM^ CD3/CD28 Human T cell Activators (STEMCELL Technologies, anti-human CD3 and anti-human CD28 monospecific antibody complexes) and expanded in 1000U/mL rhIL-2 (NCI) and split on day 4 with or without 20nM ACCi for 6 days. Human cultures were maintained in RPMI supplemented with 10% FBS, 300 mg/L L-glutamine, 2mM GlutaMAX, 100 units/mL penicillin, 100 µg/mL streptomycin, 50 µg/mL gentamycin, 25mM HEPES, 55µM 2-mercaptoethanol, and 0.2% Plasmocin mycoplasma prophylactic.

### Tumor-T Cell Transwell Coculture

5×10^5^ B16F1 tumor cells were seeded into 6-well companion plates. After 24 hours of adherence OT-1 T cells were introduced into transwell inserts (Corning) in complete T-cell media supplemented with 200 U/mL rhIL-2 (NCI) ± vehicle, ACCi (100nM), or Amidepsine A (20μM), where indicated, then harvested after 36 hours of coculture.

### Seahorse Bioanalysis

Seahorse XF Mito Stress Tests and Seahorse Real-Time ATP Rate Assays were performed using the Seahorse XFe96 Analyzer. 96-well Seahorse plates (Agilent) were coated with CellTak (Corning). T cells were plated in Seahorse XF DMEM Medium (Agilent) supplemented with 1% fetal bovine serum (FBS) and centrifuged for adherence. For Seahorse Mito Stress Test, 1µM oligomycin, 1.5uM FCCP, and 2µM rotenone/1µM antimycin A (Sigma) were injected sequentially, and the oxygen consumption rate (OCR) was measured. In some cases, 10μM etomoxir (MedChemExpress) was acutely injected prior to oligomycin. For the Seahorse Real-Time ATP Rate Assay, 1µM oligomycin and 2µM rotenone/1µM antimycin A (Sigma) were injected sequentially, and the oxygen consumption rate (OCR) and extracellular acidification rate (ECAR) were measured. In some cases, 10μM etomoxir (MedChemExpress) was acutely injected prior to oligomycin. The mitochondrial ATP production rate was quantified based on the decrease in the OCR. FCCP is used to elicit maximal respiration, however FCCP can inhibit OCR at high concentrations. FCCP dose of 1.5 μM was selected based on prior optimization during performance of Seahorse XF Cell Mito Stress test with 5 different doses of FCCP (Hurst et al., 2019; Hurst *et al*., 2020). Doses were selected based on published data (Buck et al., 2016; Hurst *et al*., 2020; van der Windt *et al*., 2012). For Seahorse normalizations, equal numbers of T cells per condition were seeded on Cell Tak-coated plates (Agilent) immediately prior to performing the assay and cell counts were verified via microscopy directly after assay performance. Spare respiratory capacity was calculated as the difference between the basal and maximal OCR readings after addition of FCCP.

### Metabolomics

*Sample Preparation:* T cells were harvested and washed with sodium chloride solution then resuspended in 80% methanol for three freeze-thaw cycles at –80°C and stored at –80°C overnight to precipitate the hydrophilic metabolites. Samples were centrifuged at 20,000xg and methanol was extracted for metabolite analysis. Remaining protein pellets were dissolved with 8M urea and total protein was quantified using Pierce BCA assay (Thermo Fisher). Total protein amount was used for equivalent loading for High Performance Liquid Chromatography and High-Resolution Mass Spectrometry and Tandem Mass Spectrometry (HPLC-MS/MS) analysis. *Data Acquisition*: Samples were dried with a SpeedVac then 50% acetonitrile was added for reconstitution followed by overtaxing for 30 sec. Sample solutions were centrifuged and supernatant was collected and analyzed by High-Performance Liquid Chromatography and High-Resolution Mass Spectrometry and Tandem Mass Spectrometry (HPLC-MS/MS). The system consists of a Thermo Q-Exactive in line with an electrospray source and an Ultimate3000 (Thermo Fisher) series HPLC consisting of a binary pump, degasser, and auto-sampler outfitted with a Xbridge Amide column (Waters; dimensions of 2.3 mm × 100 mm and a 3.5 µm particle size). The mobile phase A contained 95% (vol/vol) water, 5% (vol/vol) acetonitrile, 10 mM ammonium hydroxide, 10 mM ammonium acetate, pH = 9.0; B with 100% Acetonitrile. For gradient: 0 min, 15% A; 2.5 min, 30% A; 7 min, 43% A; 16 min, 62% A; 16.1-18 min, 75% A; 18-25 min, 15% A with flow rate of 150 μL/min. A capillary of the ESI source was set to 275 °C, with sheath gas at 35 arbitrary units, auxiliary gas at 5 arbitrary units and the spray voltage at 4.0 kV. In positive/negative polarity switching mode, an *m*/*z* scan range from 60 to 900 was chosen and MS1 data was collected at a resolution of 70,000. The automatic gain control (AGC) target was set at 1 × 10^6^ and the maximum injection time were 200 ms. The top 5 precursor ions were subsequently fragmented, in a data-dependent manner, using the higher energy collisional dissociation (HCD) cell set to 30% normalized collision energy in MS2 at a resolution power of 17,500. Besides matching m/z, metabolites were identified by matching either retention time with analytical standards and/or MS2 fragmentation pattern. Metabolite acquisition and identification was carried out by Xcalibur 4.1 software and Tracefinder 4.1 software, respectively. *Statistical Analysis:* Post identification, samples were normalized by taking the peak area under the curve for each metabolite per sample and dividing by the quotient of the total ion count per sample over the lowest total ion count in the batch. Subsequent transformation of normalized data was carried out with auto scaling to account for heteroscedasticity (van den Berg et al., 2006). Metabolites that were below detection in all samples were removed from analysis; missing values were imputed with 1/5 of the minimum positive value of their corresponding variable. Differential metabolite abundance between groups of interest was defined as absolute fold change > 2 and FDRq Adj p-value < 0.05. Quantitative enrichment analysis was performed using centered and scaled levels of all detectable metabolites after removal of invariant metabolites using the SMPDB database to identify biological processes associated with differential expression. All statistical analysis was performed using the MetaboAnalyst 5.0 web server (Pang et al., 2021).

### Lipidomics

*Sample Preparation:* Cells were harvested and washed in PBS and cell pellets were frozen at – 80°C.Pellets were extracted with a liquid/liquid partition of 250 µL water, 300 µL methanol with 1.5 µg/mL Avanti EquiSPLASH mix, and 1 mL of methyl tert-butyl ether (MTBE). Samples were shaken for 30 minutes and then centrifuged at 20,000 rcf for 10 minutes to facilitate phase separation. The organic layer was removed, dried down, and reconstituted in 100 µL of isopropyl alcohol for liquid chromatography coupled to mass spectrometry (LC-MS) analysis. *Data Acquisition*: Samples were analyzed by Ultra Performance Liquid Chromatography coupled to High-Resolution Mass Spectrometry (UPLC-MS). The system consists of a Thermo Q Exactive HF-X coupled to a Waters H Class UPLC. A 100 mm x 2.1 mm, 1.7 µm Waters BEH C18 column was used for separations. The following mobile phases were used: A-60/40 ACN/H20 B-90/10 IPA/ACN; both mobile phases had 10 mM Ammonium Formate and 0.1% Formic Acid. A flow rate of 0.2 mL/min was used. Starting composition was 32% B, which increased to 40% B at 1 min (held until 1.5 min) then 45% B at 4 minutes. This was increased to 50% B at 5 min, 60% B at 8 min, 70% B at 11 min, and 80% B at 14 min (held until 16 min). At 16 min the composition switched back to starting conditions (32% B) and was held for 4 min to re-equilibrate the column. MS^1^ data was collected at a resolution of 60,000 with an automatic gain control (AGC) target setting of 1 × 10^6^ and a maximum inject time of 50 ms in each ionization mode. Data dependent fragmentation was performed on the top 5 ions with a resolution of 15,000, an AGC target of 1× 10^5^, and a maximum inject time of 50 ms with a dynamic exclusion window of 10 sec. HCD was performed with a normalized step collision energy of 25, 35, and 45 V. Heated electrospray ionization was utilized with the following settings: sheath gas 53 units, auxiliary gas 14 units, sheath gas 2 units, spray voltage 3.5 kV (positive) and 3.5 kV (negative), capillary temperature 280°C, RF funnel 100 V, and auxiliary gas heater temperature 400°C. *Analysis:* Lipid identifications were performed in Lipid Search 4.2.2 using MS/MS fragmentation data (mass error of precursor=5 ppm, mass error of product=8 ppm). The identifications were generated individually for each sample and then aligned by grouping the samples (retention time tolerance=0.1 min). Normalization was performed using peak area of specific deuterated classes in the Avanti EquiSPLASH mix. Principal component analysis (PCA) was performed on a data matrix generated by MZmine 2.3. The following parameters were used for peak detection of the data: noise level (absolute value), 1 × 10^5^ counts; minimum peak duration 0.5 min; tolerance for *m/z* intensity variation, 20%. Peak list filtering and retention time alignment algorithms were performed to refine peak detection. The join algorithm was used to integrate all the chromatograms into a single data matrix using the following parameters: the balance between *m/z* and retention time was set at 10.0 each, m/z tolerance was set at 0.001 or 5 ppm, and retention time tolerance was defined as 0.5 min. Once an aligned peak list was created, features (*m/z* and retention time pairs) present in the first blank sample of the run were deleted as a means of blank filtering. The peak list from MZmine was exported into Excel. PCA was conducted in SIMCA 17 by Umetrics. The peak area was transformed via a fourth root transformation to reduce heteroscedastic noise.

### Proteomics

*Sample Preparation:* T cells were washed in PBS and lysed in 9M urea, 50 mM Tris pH 8, and 100 units/mL Pierce Universal Nuclease (Thermo Fisher Scientific) and the concentration of protein was measured using a Pierce BCA Assay (Thermo Fisher Scientific). Protein was trypsin (Sigma Aldrich) digested at 37°C for 18 hours, and the resulting peptides were desalted using C18 ZipTips (Millipore). *LC/MS data acquisition parameters:* Peptides were separated and analyzed on an EASY nLC 1200 System (Thermo Fisher Scientific) in-line with the Orbitrap Fusion Lumos Tribrid Mass Spectrometer (Thermo Fisher Scientific) with instrument control software v. 4.2.28.14. Peptides were pressure loaded at 1,180 bar, and separated on a C18 Reversed Phase Column [Acclaim PepMap RSLC, 75 μm x 50 cm (C18, 2 μm, 100 Å); Thermo Fisher Scientific] using a gradient of 2%–35% B in 120 minutes (Solvent A: 0.1% FA; Solvent B: 80% ACN/0.1% FA) at a flow rate of 300 nL/minute at 45°C. Mass spectra were acquired in data-dependent mode with a high resolution (60,000) FTMS survey scan, mass range of m/z 375–1,575, followed by tandem mass spectra of the most intense precursors with a cycle time of 3 seconds. The automatic gain control target value was 4.0e5 for the survey MS scan. Fragmentation was performed with a precursor isolation window of 1.6 m/z, a maximum injection time of 50 ms, and HCD collision energy of 35%; the fragments were detected in the Orbitrap at a 15,000 resolution. Monoisotopic precursor selection was set to “peptide.” Apex detection was not enabled. Precursors were dynamically excluded from resequencing for 20 seconds and a mass tolerance of 10 ppm. Advanced peak determination was not enabled. Precursor ions with charge states that were undetermined, 1, or >7 were excluded. *Mass spectrometry data processing:* Protein identification and quantification were extracted from raw LC/MS-MS data using the MaxQuant platform v.1.6.3.3 with the Andromeda database searching algorithm and label free quantification (LFQ) algorithm. Data were searched against a mouse Uniprot reference database UP000000589 with 54,425 proteins (March, 2019) and a database of common contaminants. The FDR, determined using a reversed database strategy, was set at <1% at the protein and peptide level. Fully tryptic peptides with a minimum of seven residues were required including cleavage between lysine and proline. Two missed cleavages were permitted. LC/MS-MS analyses were performed in triplicate for each biological replicate with match between runs enabled. The “fast LFQ” was disabled and “stabilize large ratios” features were enabled. The first search was performed with a 20 ppm mass tolerance, after recalibration a 4.5 ppm tolerance was used for the main search. A minimum ratio count of 2 was required for protein quantification with at least one unique peptide. Parameters included static modification of cysteine with carbamidomethyl and variable N-terminal acetylation. The protein groups’ text file from MaxQuant was processed in Perseus v. 1.6.5.0. Identified proteins were filtered to remove proteins only identified by a modified peptide, matches to the reversed database, and potential contaminants. Normalized LFQ intensities were log_2_ transformed. The LFQ intensities of technical replicates were averaged for each biological replicate. Quantitative measurements were required in at least three of five biological replicates in each treatment group. *Statistical Analysis:* Log_2_ transformed, protein LFQ intensities, normalized in MaxQuant, exhibited normal distributions. Binary comparisons of each LC/MS-MS analysis yielded Pearson correlation coefficients > 0.90 between all technical and biological replicates within each experiment. For comparison of vehicle and tumor-coculture T cells, quantitative measurements were required in at least three of five biological replicates in each group. With a permutation-based FDR of 5% and S0 = 0.01, q val < 0.05, 177 proteins were significantly different between control and tumor transwell cocultured T cells. toppgene.cchmc.org Toppfun software package was used to measure significantly enriched GO:BP pathways.

### Free Fatty Acid Analysis

*Sample preparation:* Samples were extracted with 500 µL of 2.5% sulfuric acid in methanol. Samples were heated in a 70^ₒ^ C water bath for 1 hour. Samples were allowed to cool for 5 minutes then 500 µL of water and 200 µL of hexanes were added. Samples were vortexed then centrifuged for 10 min at 20,000 rcf. The top layer, ∼150 µL, was pulled off for analysis. A standard curve was prepared in hexanes using the comprehensive fatty acid methyl ester mix from Cayman Chemical. *Analysis:* Samples were analyzed with a ThermoFisher Exactive GC (ThermoFisher, Bremen, Germany) mass spectrometer Samples were introduced via Electron Impact (EI) ionization (70 eV). The mass range was set to 50-400 m/z. All measurements were recorded at a resolution setting of 60,000. Separations were conducted on a Agilent J&W HP-88 column (100 m x 0.25 mm ID x 0.20 um film). GC conditions were set at 100°C to start and maintained for 13 min. System then heated at 10°C/min up to 180°C and was held for 6 min then to 192°C at 1°C/min and held for 9 min and then to heated to 230°C at 10°C/min and held for 10 min (total method 63 min) Injection volume for all samples was 1 uL with a 20:1 split ratio. The carrier gas (helium) was held at constant flow of 1.3 mL/min. Xcalibur was used to integrate the peaks and quantify the fatty acids.

### Microscopy

*Transmission electron microscopy*: TEM images were acquired at the Microscopy Services Laboratory at the University of North Carolina at Chapel Hill. Cells were fixed in 2% paraformaldehyde/2.5% glutaraldehyde in 0.1M sodium cacodylate and stored at 4°C until further analysis. Cells were washed and incubated in 1% osmium tetroxide in 0.15M sodium phosphate buffer for 1 hour. Cells were washed and then dehydrated through a gradient series of ethanol and two 15-minute exchanges of propylene oxide. Cell pellets were infiltrated with a 1:1 mixture of propylene oxide and Polybed 812 epoxy resin (Polysciences, Inc.) for 2 hours and infiltrated overnight. Ultrathin sections (70-80 nm) were cut using a diamond knife mounted on 200 mesh copper grids followed by staining with 4% aqueous uranyl acetate for 12 minutes and Reynold’s lead citrate for 8 minutes. Samples were observed using a JEOL JEM-1230 transmission electron microscope operating at 80kV and images were acquired with a Gatan Orius SC1000 CCD Digital Camera and Gatan Microscopy Suite 3.0 software. Quantification of mitochondrial area, cristae length, and mitochondria-ER contact lengths were performed using ImageJ software. *Confocal microscopy:* Cells were fixed in 4% paraformaldehyde in PBS for 15 minutes and permeabilized with 0.1% TritonX-100 in PBS for 15 minutes at room temperature. Cells were blocked with 2% BSA in PBS for 1 hour at room temperature before antibody staining with rabbit ⍺-Sec61b (Cell Signaling) and mouse ⍺-Tom20 (Santa Cruz) antibodies in 1% BSA in PBS for 1 hour at room temperature. Cells were washed with 1x PBS and incubated with nuclear stain DRAQ5 (Abcam), neutral lipid dye Bodipy 493/503 (Molecular Probes), and secondary antibodies goat anti-rabbit IgG (H+L) AlexaFluor 594 and goat anti-mouse IgG (H+L) AlexaFluor 488 in 1% BSA in PBS for 1 hour at room temperature. A 96-well glass bottom plate (Cellvis) was coated with CellTak (Corning), washed with DI water, air dried, and labeled cells were centrifuged for adhesion. Cells were imaged using the Zeiss LSM-880 confocal microscope with AiryScan detector and 3-D reconstruction was generated from confocal Z-stacks using Imaris software. Lipid droplets were quantified using raw images in ImageJ.

### Flow Cytometry and High Dimensional Analysis

For murine T cell phenotyping, cells were stained with Zombie NIR viability dye (Biolegend) in PBS at 4°C for 15 minutes. After wash, cells were stained with CD8-Spark Blue 550 (Biolegend), CD44-Brilliant Violet 785 (Biolegend), CD62L-BV421 (BD Biosciences), PD-1-BB700 (BD Biosciences), TIM-3-BV711 (Biolegend), LAG-3-APC-eFluor 780 (Invitrogen), ICOS-Super Bright 436 (Invitrogen), CD69-PE-Cy5 (Biolegend), CD95-BV480 (BD Biosciences), GITR-BV650 (BD Biosciences), and CD27-BV750 (BD Biosciences) for 30 minutes at 4°C. Cells were washed and fixed overnight using the Foxp3/Transcription Factor Fixation/Permeabilization Concentrate and Diluent kit and Permeabilization Buffer (eBioscience). Cells were then washed and stained with CTLA-4-PE-Dazzle 594 (Biolegend), Ki67-PerCP-eFluor 710 (Invitrogen), TOX-PE (Miltenyi Biotec), TCF1/7-PE-Cy7 (Cell Signaling Technology), Granzyme B-Alexa Fluor 700 (Biolegend), and BCL-2-Alexa Fluor 647 (Biolegend) for 3 hours. Samples were collected on a Cytek Northern Lights. For *ex vivo* analysis, TILs were restimulated with Cell Stimulation Cocktail (Invitrogen) and Brefeldin A for 4-6 hours, then collected and washed and stained with Zombie NIR viability dye (Biolegend) in PBS at 4°C for 15 minutes. After wash, cells were stained with CD8-Spark Blue 550 (Biolegend) and CD45.2-Brilliant Violet 785 (Biolegend) for 30 minutes at 4°C. Cells were washed and fixed overnight using the Foxp3/Transcription Factor Fixation/Permeabilization Concentrate and Diluent kit and Permeabilization Buffer (eBioscience). Cells were then washed and stained with TNF-α, IFN-γ for 3 hours. Samples were collected on a Cytek Northern Lights. For neutral lipid staining, cells were incubated with 1mM Bodipy^493/503^ (Life Technologies) for 30 minutes at 37C prior to staining as above. For human PBMC and T cell cultures, cells were stained with Zombie NIR viability dye (Biolegend) in PBS at 4°C for 15 minutes. After wash, cells were stained with CD8-Superbright 436 (Invitrogen), CCR7-PE (Invitrogen), CD45RA-PE-Cy7 (Invitrogen), and CD4-APC (Invitrogen). For detection of p-ACC, extracellular staining was performed as above then fixed with BD Phosflow Lyse/Fix buffer (BD) and permeabilized with perm/wash buffer 1 (BD). Cells were incubated with p-ACC antibody (Cell Signaling) then probed with goat anti-rabbit IgG-Alexa Fluor 488. Samples were collected on a Cytek Northern Lights. For all spectral flow cytometry experiments, single stain compensations were created using UltraComp eBeads Plus Compensation Beads (Invitrogen) or heat-killed cell samples. Live CD8^+^ singlets were gated and subsampled to 3000 events per sample. To facilitate visualization of marker expression patterns, cells were mapped to a two-dimensional embedding using the Uniform Manifold Approximation and Projection (UMAP) algorithm. Mean fluorescent intensities of select markers were then projected onto the UMAP space to further visualize the distinct phenotypes. All analysis was performed using the OMIQ analysis or FloJo software platforms.

### RNA-sequencing

Cells were washed in PBS and pellets were flash frozen at –80°C. RNA was extracted with a Promega Maxwell RSC 16 using the Maxwell RSC simplyRNA Cells kit according to the manufacturer’s instructions. 1 mg of total RNA was used for the construction of libraries with the New England Biolabs NEBNext® Poly(A) mRNA Magnetic Isolation Module and Ultra II Directional RNA Library Prep Kit for Illumina according to the manufacturer’s instructions. Dual-indexed libraries were pooled and sequenced at VANTAGE (Vanderbilt University Medical Center) on an Illumina NovaSeq 6000 (S4 flow cell) to a depth of approximately 25 million paired-end 150 bp reads per library. Data Processing & Analysis: Raw fastq files were then aligned to the GRCm38.p6 version of the mouse genome and transcriptome using STAR v2.7.6a. Quantification of gene abundance for each sample done by Salmon v1.4.0. Quality control evaluation of the data was done using FASTQC v0.11.9 as well as the Picard Toolkit v2.22.4 *CollectRnaSeqMetrics*, *flagstat* and *maximum read length* tools. Gene level counts were compiled, and a gene was removed from subsequent analysis if it contained fewer than 5 reads across all the samples. Principal Component Analysis and Relative Log Expression calculated to check for any outliers. Normalization and differential gene expression was performed using the DESeq2 v1.22.2 Bioconductor package in R v4.0.3. The Benjamini-Hochberg method was used to correct the *P* values for multiple hypothesis testing.

### Tumor Modeling

MCA-205 fibrosarcomas were established subcutaneously by injecting 2.5×10^5^ cells into the right flank of C57BL/6 mice. After 12-14 days of tumor growth, spleens and tumors from groups of mice were harvested and tumors were processed using the Mouse Tumor Dissociation Kit and gentleMACS dissociator (Miltenyi Biotec) according to manufacturer’s protocol. Samples were pre-enriched using EasySep Mouse CD8^+^ T Cell Isolation Kit (Stemcell Technologies) according to manufacturer’s protocol and sorted using the FACSAria II cell sorter. CD8^+^ spleen and pooled TIL samples were washed in PBS and frozen for proteomic, lipidomic, and RNA-seq analysis. For adoptive cellular therapy experiments, B16F1-OVA melanomas were established subcutaneously by injecting 2.5×10^5^ cells into the right flank of C57BL/6 mice and tumor-bearing hosts were irradiated with 5 Gy 24 hours prior to T cell transfer. After 10 days of tumor growth, 5×10^5^ OT-1 T cells conditioned with vehicle or ACCi were infused in 100 µL PBS via tail vein into tumor bearing mice. Tumor growth was measured every other day with calipers, and survival was monitored with an experimental endpoint of tumor growth ≥ 300 mm^2^. For tumor harvest experiments, 1×10^6^ vehicle or ACCi-conditioned OT-1 T cells were infused into B16-OVA tumor bearing mice 10 days after tumor challenge and tumors were harvested 7 days later using the Mouse Tumor Dissociation Kit and gentleMACS Octo dissociator with heaters (Miltenyi Biotec) according to the manufacturer’s protocol.

### Statistical Analysis

GraphPad Prism v 9.3.0 was used to calculate p values with unpaired two-sided Student’s *t-*test or paired Student’s *t-*test as indicated in figure legends. Survival analysis was calculated using Log-Rank (Mantel-Cox) test. Values of p < 0.05 were considered significant.

## Supplemental Information

Supplemental Information can be found in the Supplemental Information document.

## Acknowledgements

We are grateful to Drs. R. Luke Wiseman and Jonathan Serody for thoughtful critique of the manuscript. RNA-sequencing was performed by the MUSC Translational Science Laboratory in combination with VANTAGE at Vanderbilt University Medical Center and GSEA was performed by the Lineberger Comprehensive Cancer Center Bioinformatics Core. Proteomics was performed by the Medical University of South Carolina Proteomics Core. Metabolomics were performed by the Metabolomics Core Facility at Northwestern University. Lipidomic studies were performed by the Mass Spectrometry Core Laboratory, Department of Chemistry at UNC Chapel Hill. Transmission electron microscopy was performed by the Microscopy Services Laboratory, Department of Pathology and Laboratory Medicine at UNC Chapel Hill. Patient surgical samples were acquired through the Biorepository Shared Resource, Hollings Cancer Center, Medical University of South Carolina.

## Author Contributions

Conceptualization: E. G. Hunt and J.E. Thaxton

Methodology: E. G. Hunt, K. E. Hurst, A. S. Kennedy, B. P. Riesenberg, P. Gao, M. F. Coleman, E. D. Wallace, J. E. Thaxton

Investigation (data acquisition): E. G. Hunt, K. E. Hurst, A. S. Kennedy, B. P. Riesenberg, P. Gao, E. J. Gandy, A. M. Andrews, E. D. Wallace

Resources: P. Gao, E. D. Wallace, J.E. Thaxton

Analysis: E. G. Hunt, A. S. Kennedy, B. P. Riesenberg, M. F. Coleman, P. Gao, E. D. Wallace, J.E. Thaxton

Writing, review, and/or revision of manuscript: E. G. Hunt, K. E. Hurst, P. Gao, M. F. Coleman, E. D. Wallace, J.E. Thaxton

Visualization: E. G. Hunt, J. E. Thaxton

Project administration: K.E. Hurst, J. E. Thaxton

Supervision: K. E. Hurst, J. E. Thaxton

Funding acquisition: J. E. Thaxton

## Declaration of Interests

The authors declare no competing interests.

## Inclusion and Diversity

One or more of the authors of this manuscript self-identifies as an underrepresented ethnic minority in their geographical location. One or more of the authors of this manuscript self-identifies as a member of the LGBTQIA+ community.

## Notes

### Competing Interest Statement

The authors have declared no competing interest.

