## Supplemental Materials and Figures for "Acetyl-CoA Carboxylase Obstructs CD8^+^ T-Cell Lipid Utilization and Energy Synthesis in the Tumor Microenvironment"

**Supplemental Materials and Methods**

*Immunoblotting*

T cells were lysed in RIPA buffer (Sigma Aldrich) supplemented with Protease Inhibitor Cocktail (Cell Signaling Technology) and Phosphatase Inhibitors I and II (Sigma Aldrich). Protein concentrations were normalized using Pierce BCA Kit (Thermo Fisher Scientific) and loaded to 4%–10% agarose gels (Bio-Rad). p-ACC, ACC, DGAT1, DGAT2, β-actin, and HRP-linked anti-rabbit and mouse secondaries were obtained from Cell Signaling Technology. Phospho protein was developed with Pierce ECL Plus Western Blotting Substrate (Thermo Fisher Scientific).

*Real-time PCR*

RNA was isolated with RNeasy Mini Kit (Qiagen, 74104) and concentration was measured using the SpectraDrop Micro-Volume Microplate (Molecular Devices). Single-strand cDNA was made with 500 ng RNA using the High Capacity RNA-to-cDNA Kit (Applied Biosystems, 4387406, Thermo Fisher Scientific). Mouse TaqMan Gene Probes (Applied Biosystems, Thermo Fisher Scientific) were used to perform real-time PCR in triplicate using the QuantStudio 6 Flex Real-Time PCR System (Thermo Fisher Scientific). Gene expression for *Acaca* and *Acacb* was normalized to *Gapdh* and calculated using ΔΔCt method.

*Malonyl-CoA ELISA*

T cells were collected, washed, sonicated in PBS, and frozen overnight. Two freeze-thaw cycles were performed and lysates were pelleted to remove debris. Supernatant was collected and assayed using the Mouse malonyl-Coenzyme A ELISA Kit (Cusabio) according to manufacturer’s protocol. Samples and standards were measured at 450nm using the SpectraMax ABS Plus plate reader (Molecular Devices).

*Glucose Assay*

Medias from indicated conditions were harvested after 36 hours of tumor-T cell coculture and glucose concentrations in medias were determined using the BG 1000 Blood Glucose System on the day of transwell assay harvest.

### Supplemental Figures & Figure Legends

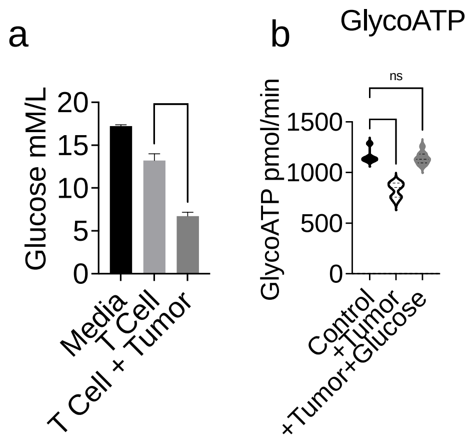


**Supplemental Figure 1.** OT-1 T cells activated with OVA peptide in the presence of IL-2 and expanded for 3 days then seeded into control or transwells seeded with B16F1 melanoma cells. After 36 hours of co-culture, control media, media from control transwells with T cells alone, or shared media from tumor cell-T cell co-cultures was collected and a) glucose concentration was measured or b) T cells were collected and Seahorse Bioanalysis Real-Time ATP Rate Assay was performed. Quantifications among T cell groups of glycolytic ATP rates (glycoATP) are shown in response to sequential injections of Oligomycin and Rotenone/Antimycin A**.** glycoATP rate was calculated as the increase in the ECAR combined with total proton efflux rate. For glucose supplementation, 25mM glucose was added to co-culture media at the time of T cell seeding. Independent experiments were repeated at least 3 times. ns, not significant, **** $p < 0.0001$ based on statistical analysis by unpaired Student's t-test, all error bars indicate the SEM. Glucose data represent average values from n=5 wells from the indicated condition. Individual Seahorse experiments repeated twice.

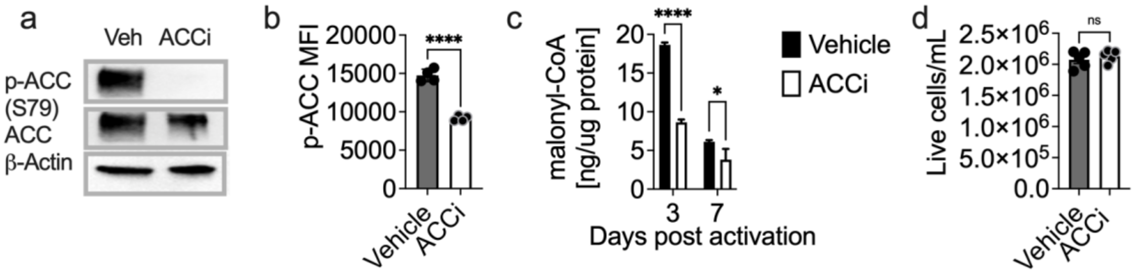


**Supplemental Figure 2. ND-646 effectively binds ACC and restricts enzymatic activity in CD8⁺ T cells.** OT-1 T cells were activated with OVA peptide in the presence of IL-2 and expanded ± ACCi. a) Immunoblot and b) FACS analysis was performed to measure p-ACC between vehicle and ACCi-treated T cell groups. c) Malonyl-CoA concentration was measured by ELISA in vehicle and therapeutic-treated T cell groups 3 and 7 days post T cell activation. d) Numbers of live OT-1 T cells/mL expanded ± ACCi for 7 days. Data points represent T cell preparations from n=5 mice. ns, not significant, * $p < 0.05$, **** $p < 0.0001$ based on statistical analysis by unpaired Student's t-test, all error bars indicate the SEM. Individual experiments performed at least twice.

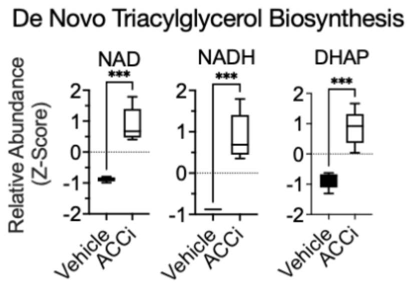


**Supplemental Figure 3.** OT-1 T cells were activated with OVA peptide in the presence of IL-2 and expanded ± ACCi for 7 days. Relative abundance of individual metabolites driving QEA signatures as in Figure 1 of *de novo* Triacylglyceride Biosynthesis between T cell preparations from 5 mice cultured ± ACCi. *** $p < 0.001$ based on statistical analysis by unpaired Student's t-test, all error bars indicate the SEM. Criteria for significance from QEA were FDR Adj $p < 0.05$.

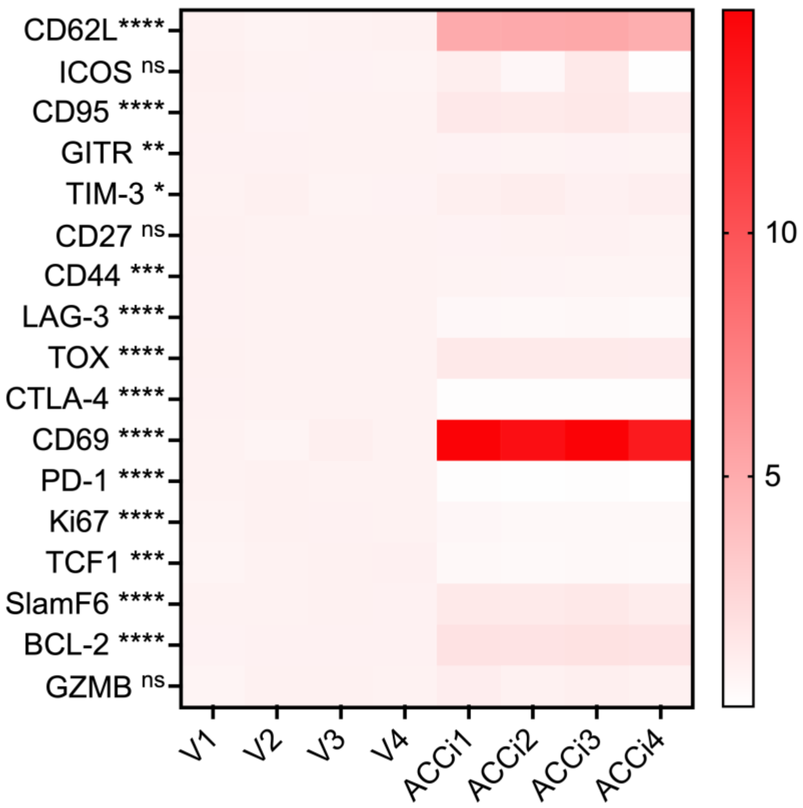


**Supplemental Figure 4.** OT-1 T cells were activated with OVA peptide in the presence of IL-2 and expanded ± ACCi for 7 days. Spectral flow cytometry was used to monitor the mean fluorescent intensities (MFI) of 20 descriptive phenotypic markers associated with T cell activation, differentiation, exhaustion, and memory. Heat map values represent scaled MFIs for a given phenotypic marker from T cell cultures generated from 4 separate mice. ns, not significant, * $p < 0.05$, ** $p < 0.01$, *** $p < 0.001$, **** $p < 0.0001$ based on statistical analysis by unpaired Student's t-test. Individual experiment repeated at least 3 times.
